# SARS-Cov-2 Spike binding to ACE2 is stronger and longer ranged due to glycan interaction

**DOI:** 10.1101/2021.07.15.452507

**Authors:** Yihan Huang, Bradley S. Harris, Shiaki A. Minami, Seongwon Jung, Priya S. Shah, Somen Nandi, Karen A. McDonald, Roland Faller

## Abstract

Highly detailed steered Molecular Dynamics simulations are performed on differently glycosylated receptor binding domains of the SARS-CoV-2 spike protein. The binding strength and the binding range increases with glycosylation. The interaction energy rises very quickly with pulling the proteins apart and only slowly drops at larger distances. We see a catch slip type behavior where interactions during pulling break and are taken over by new interactions forming. The dominant interaction mode are hydrogen bonds but Lennard-Jones and electrostatic interactions are relevant as well.

**Statement of Significance:** Glycosylation of the receptor binding domain of the Spike protein of SARS-CoV-2 as well as the ACE2 receptor leads to stronger and longer ranged binding interactions between the proteins. Particularly, at shorter distances the interactions are between residues of the proteins themselves whereas at larger distances these interactions are mediated by the glycans.

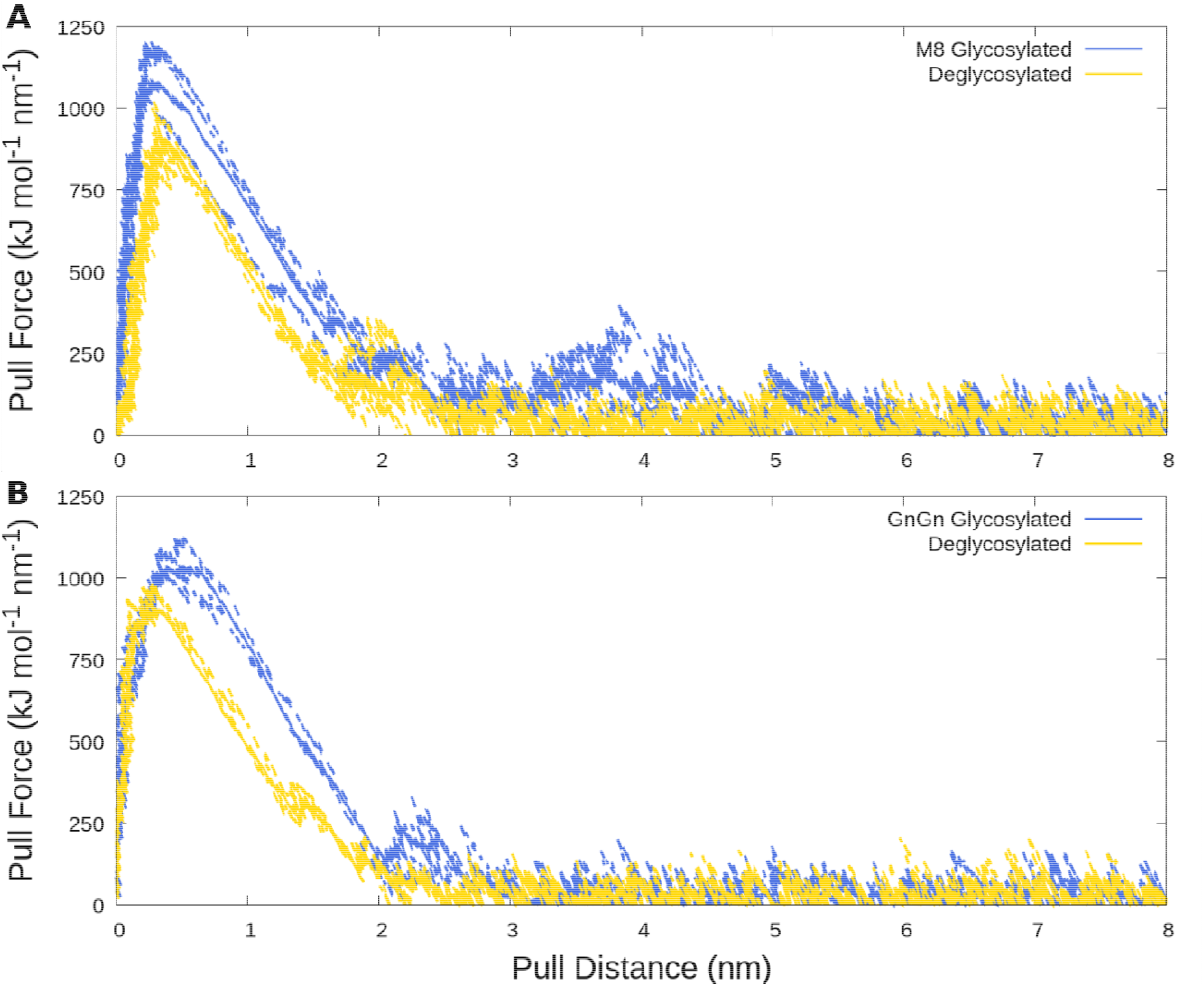

## Introduction

As of July 2021, more than 182 million people globally have been confirmed to be infected with severe acute respiratory syndrome coronavirus 2 (SARS-CoV-2), which causes coronavirus infectious disease 2019 (COVID-19). This zoonotic pandemic has disrupted society and spurred a wide range of scientific endeavors to improve our knowledge of coronaviruses and address the crisis. As the disease spreads and in order to prepare for potential future events there is a critical need for understanding the interaction of the virus with proteins involved in infection and immune clearance, or with proteins used as potential countermeasures or for the purpose of improved tests. Here, we study the interactions between SARS-CoV-2 and the human receptor responsible for binding using a molecular dynamics approach and validate it experimentally.

The SARS-CoV-2 spike (S) protein is a major structural protein and is therefore involved in many interactions. Through the receptor binding domain (RBD), S binds to the human angiotensin converting enzyme 2 (hACE2 or ACE2) receptor on the cell surface and initiates infection. There has been significant effort directed at understanding this interaction both experimentally and computationally (1–7). Such studies are critical for the development of more efficient tests and therapeutics including vaccines.

Viral structural proteins like S are often glycosylated to help pathogens evade the host immune system, modulate access to proteases, and enhance the cellular attachment through modification of protein structure and/or direct participation at the virus-host interface (8–14). Furthermore, many mammalian viruses use glycans on cell-surface glycoproteins or glycolipids as receptors (15). Despite the important role of glycans in virus-host interactions, the glycans themselves are often only partially resolved in experimental structures generated from experimental techniques such as CryoEM (16). Computational modeling of these glycans is therefore helpful in predicting their behavior and structural contributions.

S is a trimer where each monomer is expected to be highly glycosylated with 22 N-linked glycosylation sequons and 4 O-linked predicted glycosylation sites (17). Only 16 N-linked glycosylation sites were observed in a cryo-EM map of S produced in HEK293F cells (18). A study by Watanabe *et al*. (2020) determined site-specific glycoform analysis of full-length trimeric S protein made in HEK293F cells (16). In another study of S glycosylation patterns including O glycosylation were determined (19). In a similar vein, it has recently been argued that glycosylation can have influences post-vaccination and for vaccine resistance (20). Yet, the influence of glycosylation on the S-ACE2 interaction has been studied to a lesser extent (21,22). We address this gap in knowledge in the current study to reveal how glycans modulate the interaction of S with ACE2.

We expect that, as both S and ACE2 are glycosylated, the interaction is possibly modulated by the glycans. Few computational studies explicitly take the glycosylation of the receptor and/or the virus into account (23–26). This is true in general as glycosylation has only very recently become a stronger focus in simulations (27–31). One previous study has addressed the free energy of binding between the RBD and ACE2, including the impact of protein glycosylation (32). However, previous studies were limited to a single simple glycan model, and did not study interactions of glycans or the influence of different complex glycan distributions beyond pulling force and protein contacts. Additional studies have shown experimentally and computationally that the RBD and ACE2 have different binding strength and dissociation rates when they are glycosylated vs non-glycosylated (33,34). However, previous computational efforts often used simpler models for the glycans. We earlier developed a fully glycosylated model for the SARS-CoV-2 RBD and ACE2 proteins with different glycosylation patterns (2). We extend this model here to explore how a combination of complex glycans impact the energy and duration of binding. This is particularly important to improve rapid tests where viral antigens may be made in a variety of hosts with different glycan distributions.

In our previous study, we modeled ACE2 combined with the Fc domain as a therapeutic decoy. The extracellular domain of ACE2 was fused with the Fc region of human immunoglobulin, IgG1 (7). The fusion ACE2 to the Fc domain of IgG1 has several advantages as a therapeutic decoy since it increases circulatory half-life and facilitates purification through the use of the common Protein A affinity chromatography platform. This served to neutralize the S protein on the virus and block the S protein’s binding to cellular ACE2 for virus entry. ACE2-Fc was also modeled with plant glycosylation patterns. Due to the anticipated demand for high-speed production of the recombinant ACE2-Fc, plant-based transient expression systems are well-suited for rapid production. Plant cells can readily produce glycoproteins with either native, plant glycosylation (35) or with modified human-like glycoforms through genetic manipulation (36). We simulated two plant glycovariants of ACE2-Fc in our previous work: Variant 1 was targeted for ER retention with high mannose glycoforms, and Variant 2 was targeted for secretion with plant complex glycoforms. Since heterologous glycoproteins can be retained in the ER by adding a C-terminal H/KDEL-tag and the formation of Man8GlcNAc2 (Man8) N-glycans is typical for H/KDEL-tagging (37), Variant 1 was fully glycosylated with MAN8 glycans. Variant 2 was fully glycosylated with GlcNAc2XylFucMan3GlcNAc2 (GnGnXF^3^) that is a standard plant glycoform, and the S protein fragment was glycosylated with ANaF^6^ (2). Figure 1 shows the glycans used in our systems. In our previous study we simulated the influence of the two glycoforms on the interaction of S protein and the specific recombinant ACE2-Fc fusion protein. We expect that the glycosylation influence is not restricted to the fusion proteins. In this study we focus on the contribution of these different glycosylation patterns on the protein-protein interactions via hydrogen bonding, interaction energies, and determine the corresponding free energies.

**Figure 1:**
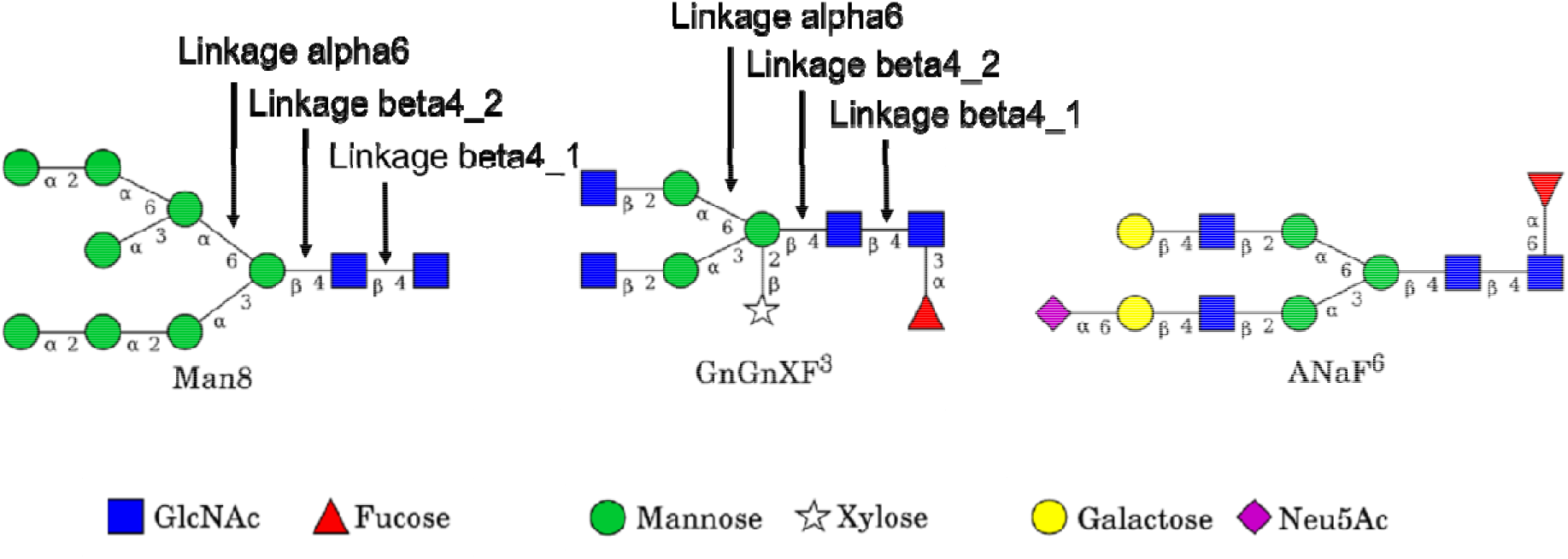
Glycans used in the simulations, adapted from previous work (2), with linkages of interest in MAN8 and GnGnXF^3^ glycans for dynamic analysis.

## Materials and Methods

### Simulation

Binding between the receptor binding domain of spike (RBD) and ACE2 receptor was determined using steered molecular dynamics, also known as the pulling of proteins (38). The starting atomic coordinates for all pulling systems were taken from the final 75 ns configurations of our previous paper (2). In that paper two sequence variants of ACE2-Fc were used to model the interaction between ACE2-F_C_ and SARS-CoV-2 RBD. Variant 1 (AF^M8^/SpFr) contained a C-terminal SEKDEL tag which is used for ER retained proteins to express high mannose glycoforms and Variant 2 (AF^GG^/SpFr) which does not contain the SEKDEL tag and expresses standard plant glycoforms. ACE2-B0AT1 and ACE2-B0AT1/SpFr structures were obtained from the protein data base. These structures had been determined using cryo-electron microscopy (PDB codes 6M18 and 6M17 (39)). These structures were fused to the Fc domain (PDB 3SGJ (40)). The Zn^2+^ coordinating residues and water were taken from structure PDB 1R42 (41) in the case of Variant 1 ACE2. Variant 2 has 2 mutations that prevent Zn^2+^ coordination. The presence of zinc in protein structures is still actively being studied to determine its role in adjusting binding specificity (42,43). It has been demonstrated that Zn^2+^ plays a role in stabilizing some protein structures and can aid in the formation of biological oligomers (42,43). The final frame of the 75 ns trajectories for both ACE2-Fc/SpFr Variants was selected, and proteins were trimmed at residue 780 ALA (Figure 2) to make the pulling simulations a manageable 851 residues with glycans and 780 residues without glycans for AF^M8^/SpFr, and 845 residues with glycans and 780 residues without glycans for AF^GG^/SpFr. Because the system changed, the force field files had to be regenerated using AmberTools (44) as described previously (2). Briefly, the molecules were trimmed and glycans were removed, then Man8 glycans were reattached to the truncated Variant 1 of ACE2, GnGnXF^3^ to the truncated Variant 2 of ACE2 and ANaF^6^ to the SpFr in both variants using Glycam.org (45). The coordinating Zn^2+^ was reattached to truncated and glycosylated Variant 1 using MCPB.py (46). Special care was taken to align the shortened original coordinates and the newly generated force field. Truncations from Variant 1 and Variant 2 that remained aglycosylated for both ACE2 and RBD were also studied to compare the influence of glycosylation on binding. The truncated systems were named A1Fr^M8^/SpFr, A1Fr/SpFr, A2Fr^GG^/SpFr, and A2Fr/SpFr, respectively. All amino acid sequences are available in supporting information and all S-S bridges are retained in our simulations.

**Figure 2:**
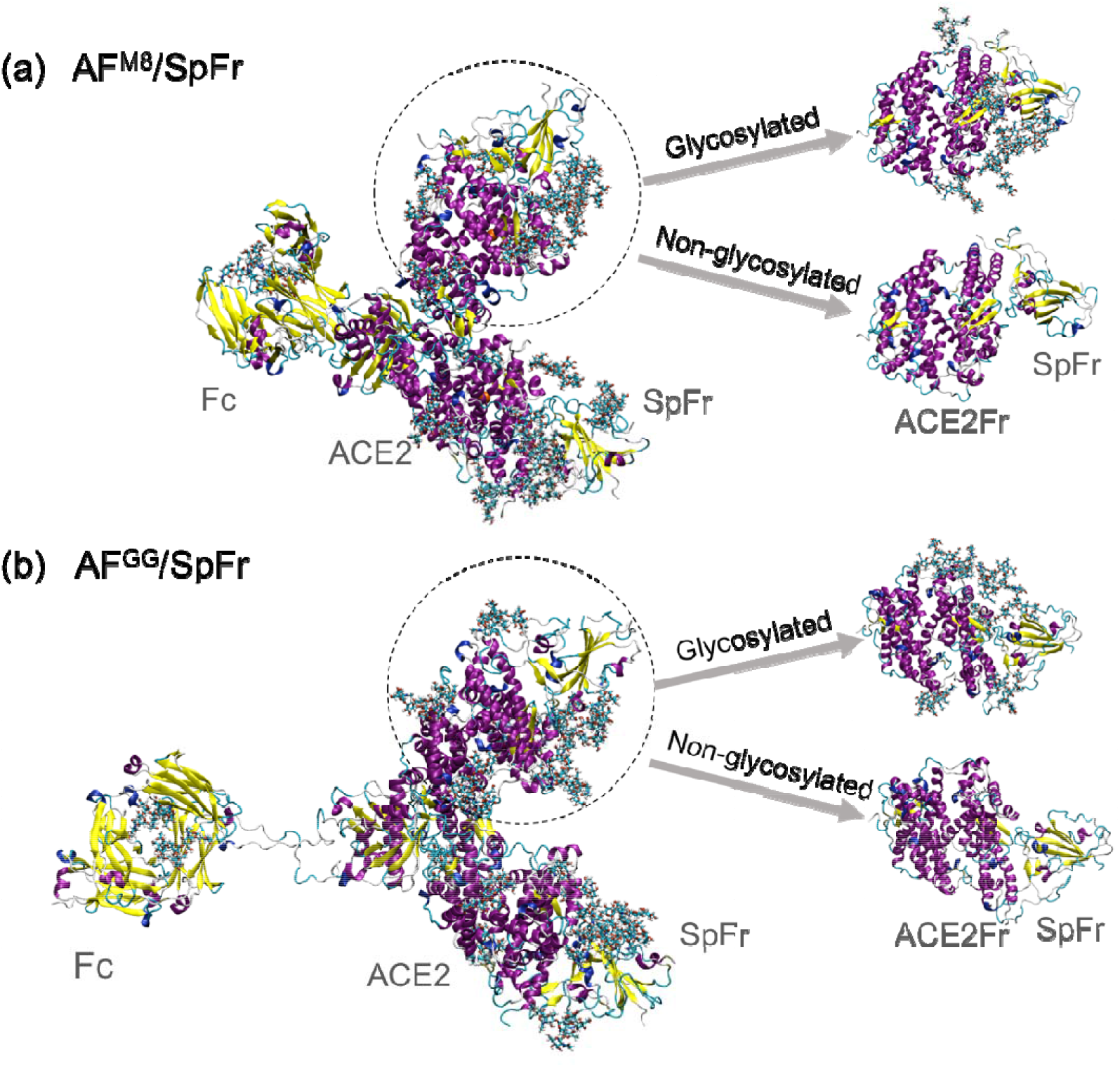
Schematics of generating the different initial structures. The full recombinantly made Ace2-FC systems on the left are used in the BLI experiments for determining binding affinities to the RBD, whereas the four truncated systems on the right containing only a fragment of ACE2 are modeled in simulations. From top to bottom, the truncated systems correspond to A1Fr^M8^/SpFr, A1Fr/SpFr, A2Fr^GG^/SpFr, and A2Fr/SpFr, respectively.

After the initial structures and corresponding force fields were generated, the proteins were rotated so that the pulling direction was along one of the principal axes, and the simulation boxes were expanded to 10 × 10 × 26 nm for A1Fr^M8^/SpFr and A1Fr/SpFr, and 10 × 10 × 30 nm for A2Fr^GG^/SpFr and A2Fr/SpFr so that the spike RBD fragments did not experience interactions with the ACE2 fragments across periodic boundaries during pulling. Then the new box was solvated with 80,271 water molecules and 24 Na^+^ as counter ions for A1Fr^M8^/SpFr, 80,764 waters and 23 Na^+^ cations for A1Fr/SpFr, 93,541 waters and 26 Na^+^ cations for A2Fr^GG^/SpFr, 93,989 waters and 25 Na^+^ cations for A2Fr/SpFr. Energy minimizations were performed until the convergence criteria were met (emtol = 1,000 kJ/mol/nm), followed by a 100 ps constant volume (NVT) (dt = 2 fs, T = 310 K) and a 100 ps constant pressure (NPT) (dt = 2 fs, T = 310 K, P = 1 atm), to equilibrate the systems. All simulations for equilibration were performed at 310 K and 1 atm with the Velocity Rescale thermostat (47) and Parrinello-Rahman barostat (48). All water bonds were constrained with SETTLE (49), and all other bonds were constrained with LINCS (50). Box expansion, solvation, and equilibration were performed using the Gromacs suite version 2019.1 (51).

Pulling simulations were then performed to study the free energy of binding as well as the structural arrangement of the separating proteins during interaction. For both variants, the ACE2 fragment was set to be immobile but deformable, whereas the spike RBD fragment (also flexible) was pulled away from the ACE2 fragment. Pull simulations were performed under NpT conditions using a 2 fs timestep, a pull coordination spring constant of 1000 kJ/mol/nm^2^, a Nose-Hoover thermostat (52) at 310 K, and a Parrinello-Rahman barostat at 1 atm.

A total of 36 pulling simulations were performed at three different pulling rates (1 nm/ns, 5 nm/ns and 10 nm/ns) on the four truncated structures using Gromacs 2019.1 (51). Each structure was pulled at each rate 3 times for sampling purposes. The starting configuration was the same for each independent run, but the random seed for the velocities in each run was randomly assigned, resulting in independent behaviors. This approach clearly generated independent runs as seen in Figure 3. Systems were pulled over a distance of 8 nm until full separation (no interaction) was achieved (see Figure 3).

**Figure 3:**
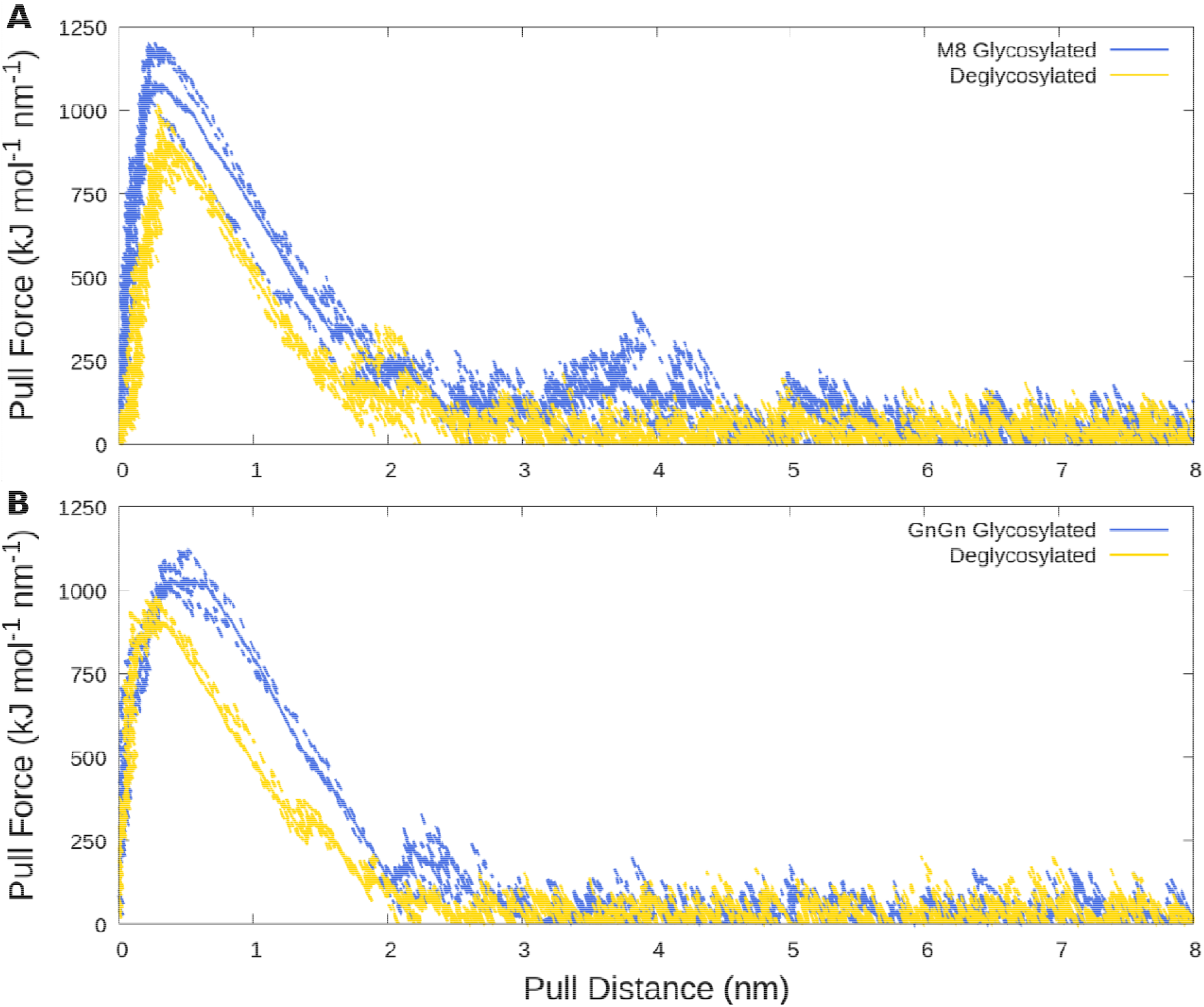
Traces of pull force versus pull distance. A) Man8 glycosylated A1Fr^M8^/SpFr and aglycosylated structure A1Fr/SpFr B) GnGnXF glycosylated A2Fr^GG^/SpFr. andaglycosylated Structure A2Fr/SpFr. Blue lines correspond to glycosylated structures, gold to deglycosylated. Dashed lines are individual replicas, solid lines are averages.

Hydrogen bonds were analyzed using the built-in Gromacs bond command (51) with a default cutoff distance of 3.5 Angstroms. This command was used to generate the hydrogen bonds and Lennard Jones contacts as a function of time as well as a hydrogen bond interaction bitmap and corresponding index file of the different interactions. The hydrogen bonding interaction bitmap was recreated in python using matplotlib (53) in order to add labels for donor acceptor pairs and calculate the percent occupancy of hydrogen bonds across the simulation (script information available in SI). Short range Lennard Jones and Coulombic interaction energies were calculated from the Gromacs. edr file by specifying energy groups on the ACE2 and RBD using the gmx energy command (54).

### Experiments

#### Protein Deglycosylation

ACE2-Fc (Acro Biosystems, Newark, DE, AC2-H5257) and RBD (Sino Biological, Chesterbrook, PA, 40592-V08B) deglycosylation was performed using Remove-iT PNGase F (Bio-Rad, Hercules, CA). Samples with PNGase F were incubated at 310 K for 5 hours. PNGase F was then removed by incubating the samples in chitin magnetic beads according to manufacturer instructions (New England Biolabs, Ipswich, MA). Deglycosylation of proteins was confirmed via sodium dodecyl sulfate polyacrylamide gel electrophoresis (SDS-PAGE). 8 μL of Laemmli sample buffer (Bio-Rad) and 2 μL β-mercaptoethanol (Bio-Rad) were added to 30 μL of sample. Samples were heated at 368 K for 5 minutes, then run on Mini-PROTEAN TGX Stain-Free Precast Gels (Bio-Rad) at 200V for 36 minutes. Gels were imaged using a ChemiDoc Imaging System (Bio-Rad).

#### Biolayer Interferometry

Anti-hIgG-Fc (AHC) biosensors (FortéBio, Fremont, CA) were used to immobilize ACE2-Fc by immersing the biosensors in solution containing 100 nM ACE2-Fc for 10 minutes. The Octet RED384 was used to obtain response measurements for protein association and dissociation. Two-fold serial dilutions of RBD were tested, from 250 nM to 7.81 nM. Data were collected for 60 seconds for the baseline, 400 seconds for association, and 800 seconds for dissociation. The experiment was performed at 299 K.

FortéBio Data Analysis Software version 8.1.0.53 was used for data processing and analysis. From the raw data, reference well values were subtracted, the y-axes were aligned to baseline, inter-step correction was applied for alignment to dissociation, and Savitzky-Golay Filtering (55) was used for smoothing. Using a 1:1 binding model, steady-state analysis was performed on the response average from 390-395 seconds. From the binding affinities of glycosylated and deglycosylated ACE2-Fc and RBD, the change in binding energy following deglycosylation of ACE2-Fc and RBD was calculated as:

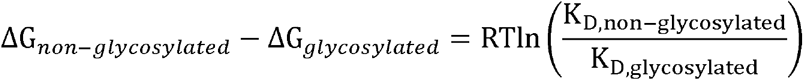

## Results

Figure 3 presents the pull force as a function of the pull distance between the ACE2 fragments and RBD for different glycosylation states at 1 nm/ns pulling rate. The pull distances are calculated based on the centers of mass for the ACE2 fragments and RBD but normalized to start from 0 nm to highlight differences between configurations. Pull force vs pull distance plots for higher pulling rates can be seen in supplemental information (Figure S1). Fundamentally, we see that for all conditions under study there is an immediate sharp increase in force when pulling the two proteins away from each other indicating strong local binding between the ACE2 binding domain and RBD. After going through a peak in force, the force drops off at increasing distance but with a clearly smaller slope than the initial increase. As expected, the pull force increases with pulling rate (blue, orange, green lines in Figure S1) such that the lowest force is most relevant for comparison to experiments. Importantly, for the same fragment the peak force is clearly higher by ~250 kJ /mol /nm at 1nm/ns, with glycosylation than without. This indicates an overall stronger binding of the glycoproteins than their aglycosylated counterparts for both types of glycosylation simulated. Additionally, the force curves are much broader for the glycosylated structures as compared to the aglycosylated ones indicating the presence of glycans extends the range for binding in addition to strengthening it. Also, the force is longer ranged (only at larger distances does it reach zero) which indicates that the glycans which extend away from the proteins contribute to the binding at longer distances. As shown in Figures 3a and 3b the aglycosylated structures return to baseline at roughly 2.5 nm of pulling distance. Importantly the glycosylated structures in Figure 3a and 3b have an extended window of pulling force of 2-3 nm for A1Fr^M8^/SpFr, and a smaller difference of roughly 1 nm for A1Fr^GG^/SpFr when compared to their aglycosylated counterparts. This indicates both Man8 and GnGnXF glycans increase binding strength, and binding range, but the type of glycan affects both the strength and interaction distance of the specific binding.

To further characterize the extension of binding interactions, Figure 4 shows hydrogen bonding interaction maps between the ACE2 and RBD proteins. Figure 4a and 4c are for A1Fr^M8^/SpFr and A2Fr^GG^/SpFr respectively while 4b and 4d are the corresponding aglycosylated versions. (Full scale images with donor:acceptor pairs labeled are available in Figures S2-S5) The y-axis contains information about the donor and acceptor pair for the hydrogen bond and the x-axis corresponds to simulation time. Interaction types are colored and sorted according to the interaction type: proteinprotein interactions are colored as white, protein-glycan as yellow, and glycan-glycan as magenta. Hydrogen bonding is clearly a major interaction mode between proteins. It is interesting that in A1Fr^M8^/SpFr (Figure 4a) the predominant interactions involve glycans directly while for A2Fr^GG^/SpFr (Figure 4c) the predominant interactions are proteinprotein interactions which are indirectly strengthened by glycosylation. This indirect protein-protein strengthening is most clearly seen when comparing occupancy calculated from these heatmaps as shown in the tables in Figure 5 and Figures S6-9. There are multiple binding regimes as a function of time for the two glycosylated structures; this is more pronounced in the A1Fr^M8^/SpFr case. This behavior manifests itself due to the original active hydrogen bonds in the complex releasing, but other hydrogen bonds catch and eventually release at larger distances before complete unbinding is seen. This catch-slip behavior is particularly attributable to the glycans, as the H-bonds present at longer distance are particularly ones involving glycans, either protein-glycan or direct glycan-glycan bonding. Both non-glycosylated structures shown in Figure 4b and 4d express maps of similar protein-protein interactions, though the A2Fr/SpFr shown in Figure 4d contains many more interactions as indicated by the increased number of rows.

**Figure 4:**
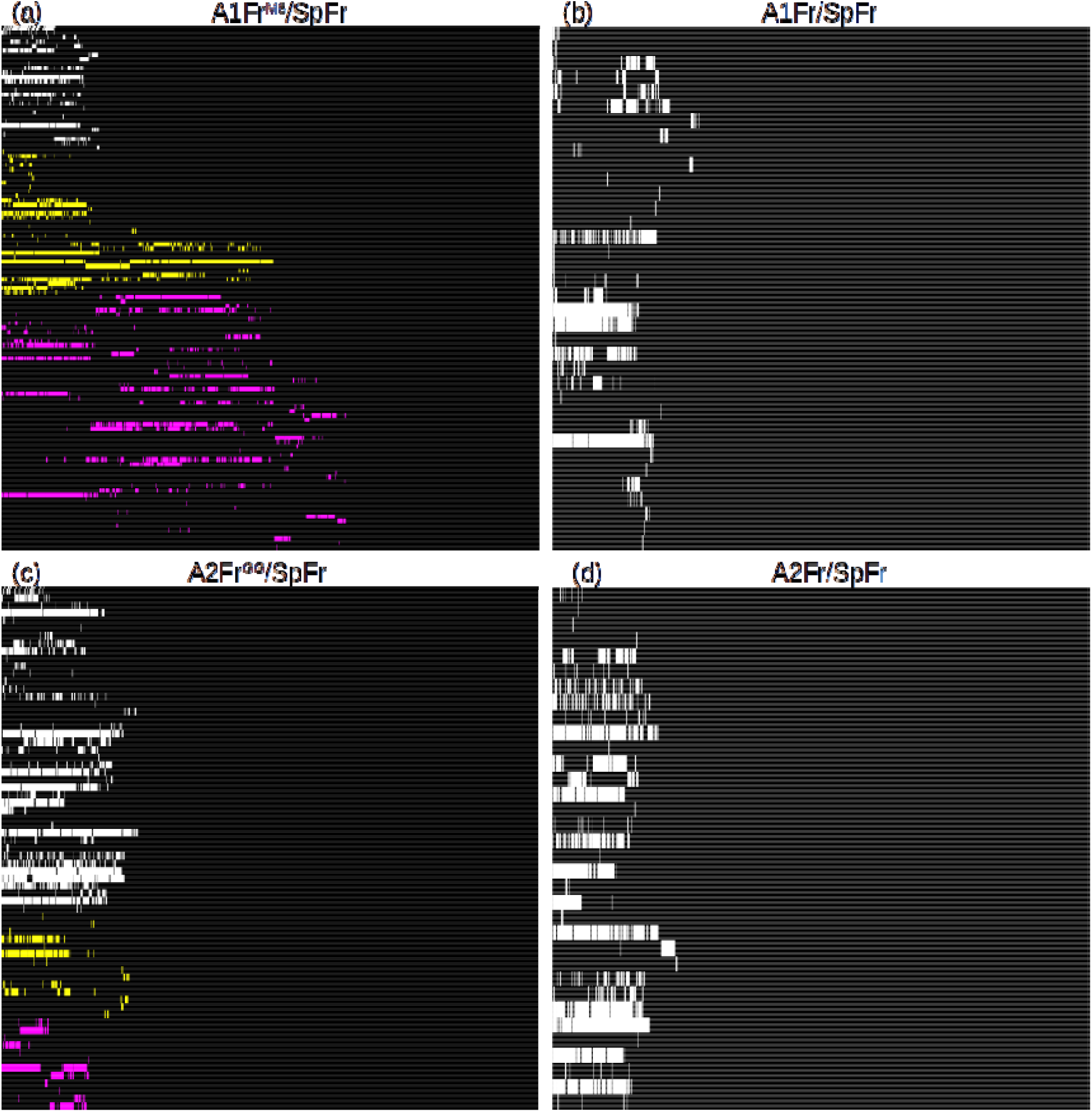
Hydrogen bond interactions vs simulation time. A) Man8 Glycosylated A1Fr^M8^/SpFr. B) Aglycosylated Structure A1Fr/SpFr. C) GnGnXF^3^ Glycosylated A2Fr^GG^/SpFr. D) Aglycosylated Structure A2Fr/SpFr. Colors indicate interaction type: White: protein-protein, Yellow: protein-glycan, Magenta: glycan-glycan.

**Figure 5:**
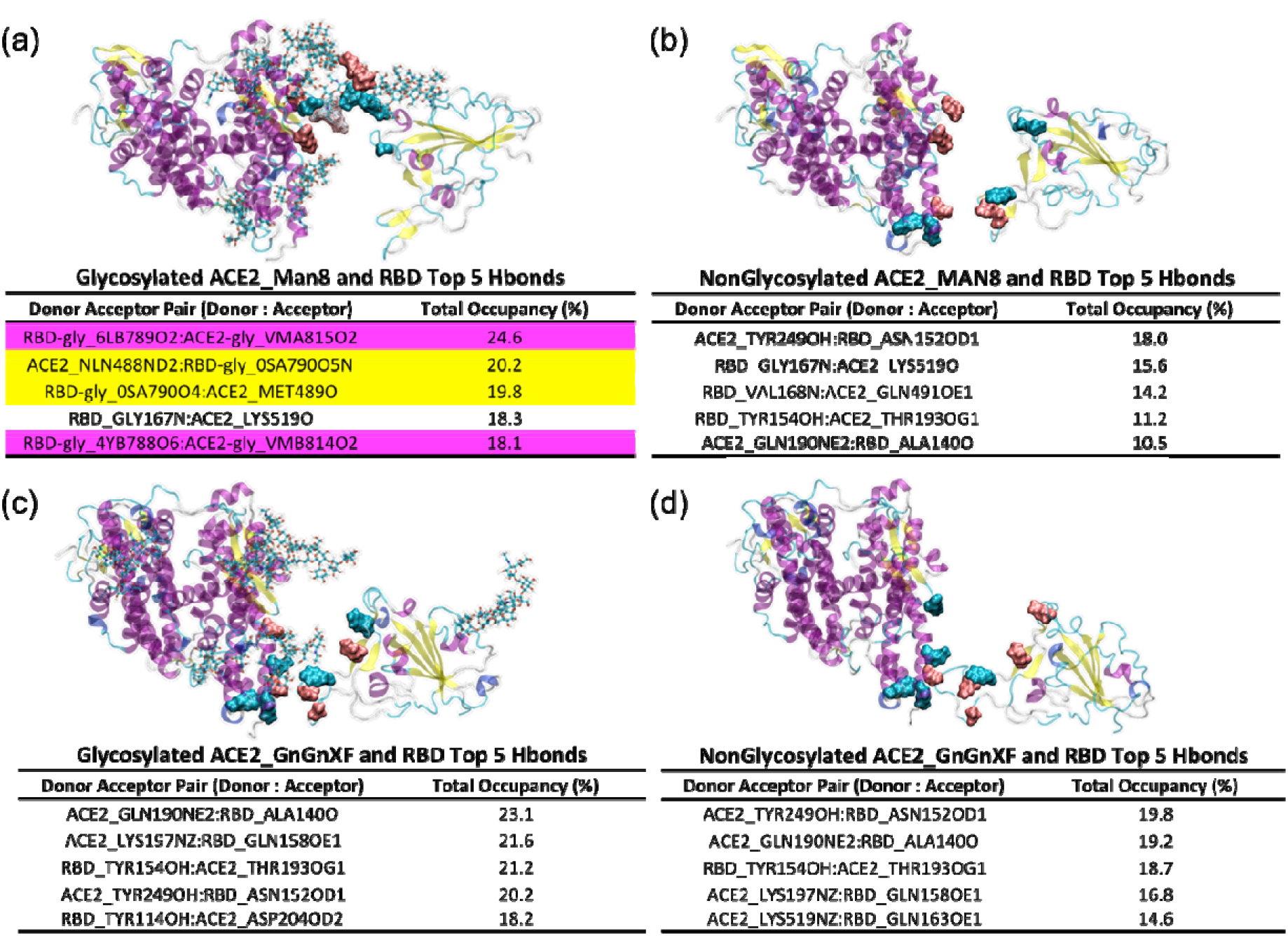
Top 5 hydrogen bond donor:acceptor pairs and occupancy. A) Man8 Glycosylated A1Fr^M8^/SpFr. B) Aglycosylated Structure A1Fr/SpFr. C) GnGnXF^3^ Glycosylated A2Fr^GG^/SpFr. D) Aglycosylated Structure A2Fr/SpFr. Table colors indicate interaction type: White: protein-protein, Yellow: protein-glycan, Magenta: glycan-glycan. On the 4 configurations, residues highlighted with blue indicate donors, and pink indicate acceptors.

Figure 5 shows the configurations where RBD with and without ANaF^6^ started to be pulled away from the ACE2 fragment for the 4 different systems. The top 5 hydrogen bonds by occupancy, i.e. the fraction of time a given hydrogen bond is active, and their corresponding donor:acceptor pairs are highlighted. (Top 25 hydrogen bonds by occupancy for the 4 different configurations are available in Figures S6-S9) A1Fr^M8^/SpFr clearly shows the predominant interactions are between the RBD glycan and ACE2 glycan and between the RBD glycan and the ACE2 protein, while for A2Fr^GG^/SpFr the predominant interactions are between the protein backbones. It is also interesting to note that the predominant interactions in A2Fr^GG^/SpFr are the protein-protein interactions. The strongest glycan interaction for A2Fr^GG^/SpFr are not found until hydrogen bond #9 ranked by occupancy (Figure S8) while the top 3 hydrogen bonds ranked by occupancy involve glycans for A1Fr^M8^/SpFr. A1Fr^M8^/SpFr also clearly shows a different starting orientation than A2Fr^GG^/SpFr, with minor changes in ACE2 structure and obvious rotation in the RBD with direct glycan-glycan interaction. These minor structural and orientational differences are also seen in the aglycosylated structures. Interacting groups for the hydrogen bonding shown follow AMBER nomenclature (56). The first letter corresponds to element with subsequent letters and numbers being linkage bookkeeping. For example, N, NZ, and NE2 all refer to nitrogen with different linkages, while O and its variants refer to Oxygen.

Figure 6 shows how the different structures of MAN8 and GnGnXF^3^ affect the hydrogen bonding regime. Although MAN8 and GnGnXF^3^ have similar size (223 atoms vs 222 atoms), their shapes are very different. MAN8 is relatively flatter comparing to GnGnXF^3^, making it bend less flexibly. Therefore, when MAN8 is close to ANaF^6^, they interact in a side-by-side fashion, whereas when GnGnXF^3^ is close to ANaF^6^, they interact in a head-to-head fashion, forming less hydrogen bonds than the MAN8/GnGnXF^3^ pair.

**Figure 6:**
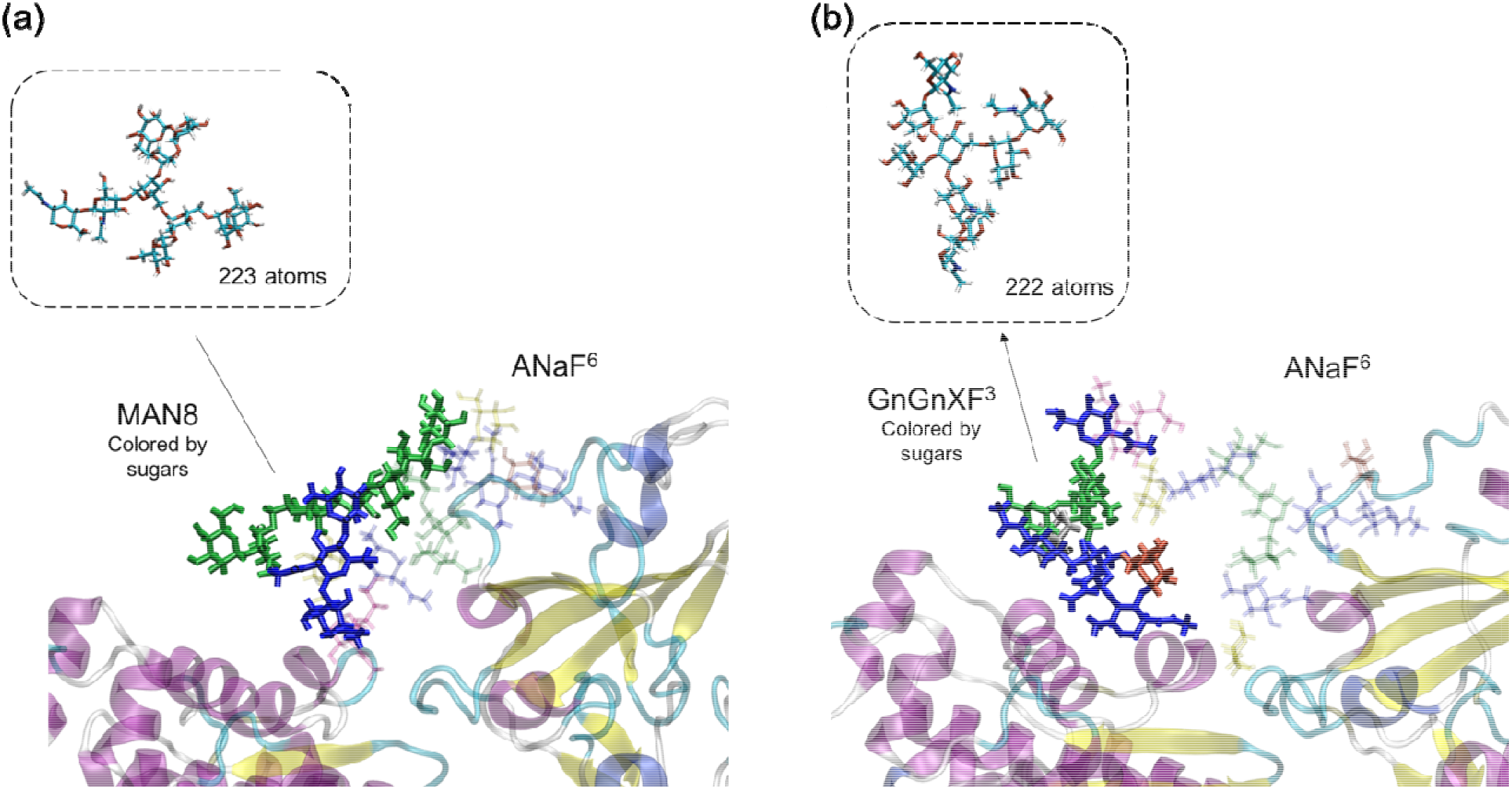
Different structures and hydrogen bonding regimes of MAN8 and GnGnXF^3^ when interacting with ANaF^6^ on RBD. A) MAN8 that interacts with ANaF^6^. B) GnGnXF^3^ that interacts with ANaF^6^. Inserts: shape and size of the MAN8 and GnGnXF^3^ without bending towards ANaF^6^. The glycans attached to proteins were colored by different sugars: Blue: GlcNAc; Green: Mannose; Yellow: Galactose; Red: Fucose; Silver: Xylose; Purple: Neu5Ac.

An autocorrelation function (ACF) analysis was performed for the angles and dihedrals of interest in both glycosylations, MAN8 and GnGnXF^3^, to further study the flexibility of the different glycans. These different flexibilities might be able to explain some of the emerging hydrogen bonding patterns. The angles and dihedrals chosen for the analysis are the ones between sugars, i.e., at the linkages. Figure 1 shows the linkages of interest; the angles and dihedrals at linkage beta4_1, beta4_2, and alpha6 of the glycans at the 6 glycosylation sites on the ACE2 fragment in A1Fr^M8^/SpFr and A2Fr^GG^/SpFr at positions N219, N256, N269, N488, N598, N712, were studied. We specifically focused on glycans at N488 for both systems as it interacts with ANaF^6^ on RBD. To improve statistics, trajectories from the previous 75 ns runs (2) were used for the ACF analysis. Figure 7 shows the angle and dihedral motions for both MAN8 and GnGnXF^3^ at glycosylation sites N219, N269, and N488. ACF results for glycans at all 6 sites are available in Figures S10, S11. Glycans on sites N219 and N269 show typical ACF behaviors of all glycans that do not directly interact with ANaF^6^ on RBD. Comparing the angle motion with dihedral motion for both glycans, ACF _Angle_ decreases significantly whereas ACF _Dihedral_ decrease slowly, indicating that angle motions are more favored for glycans and dihedral motions are constrained (alpha6 at N269 in MAN8 is the only exception where two motions are similarly favored). Comparing ACF of the different linkages, ACF of linkage alpha6 decreases much faster than the 2 beta4 linkages, indicating that linkage alpha6, which is the linkage to the branches, is the most flexible linkage. Comparing ACF of MAN8 and GnGnXF^3^, ACF _Angle_ and ACF _Dihedral_ of MAN8 decrease either at similar rate or slower than those of GnGnXF^3^ with very few exceptions (angle: N219_beta4-2, N598_beta4-2 (Figure S10); dihedral: N219_alpha6, N256_beta4-1 (Figure S11), indicating that MAN8 is generally less flexible than GnGnXF^3^ for the angle and dihedral motions at linkage beta4_1, beta4_2, and alpha6. The glycans at N488 are the ones interacting with ANaF^6^ on RBD. All angle motions and dihedral motions of MAN8 at N488 are less flexible than for GnGnXF^3^, which further proves that side-by-side hydrogen bonding fashion with ANaF^6^ is favored by MAN8 resulting in more hydrogen bonds between glycans before pulling, whereas a head-to-head arrangement is favored by GnGnXF^3^ resulting in less hydrogen bonds between glycans before pulling. In addition, the angle motions of glycans at N488 are more constrained than those of glycans at N219, and the dihedral motion of glycans at N488 are more constrained than those of glycans at N269, indicating that glycans at N488 are generally constrained because they are connected to the protein on one end, and interacting with ANaF^6^ on the other end.

**Figure 7:**
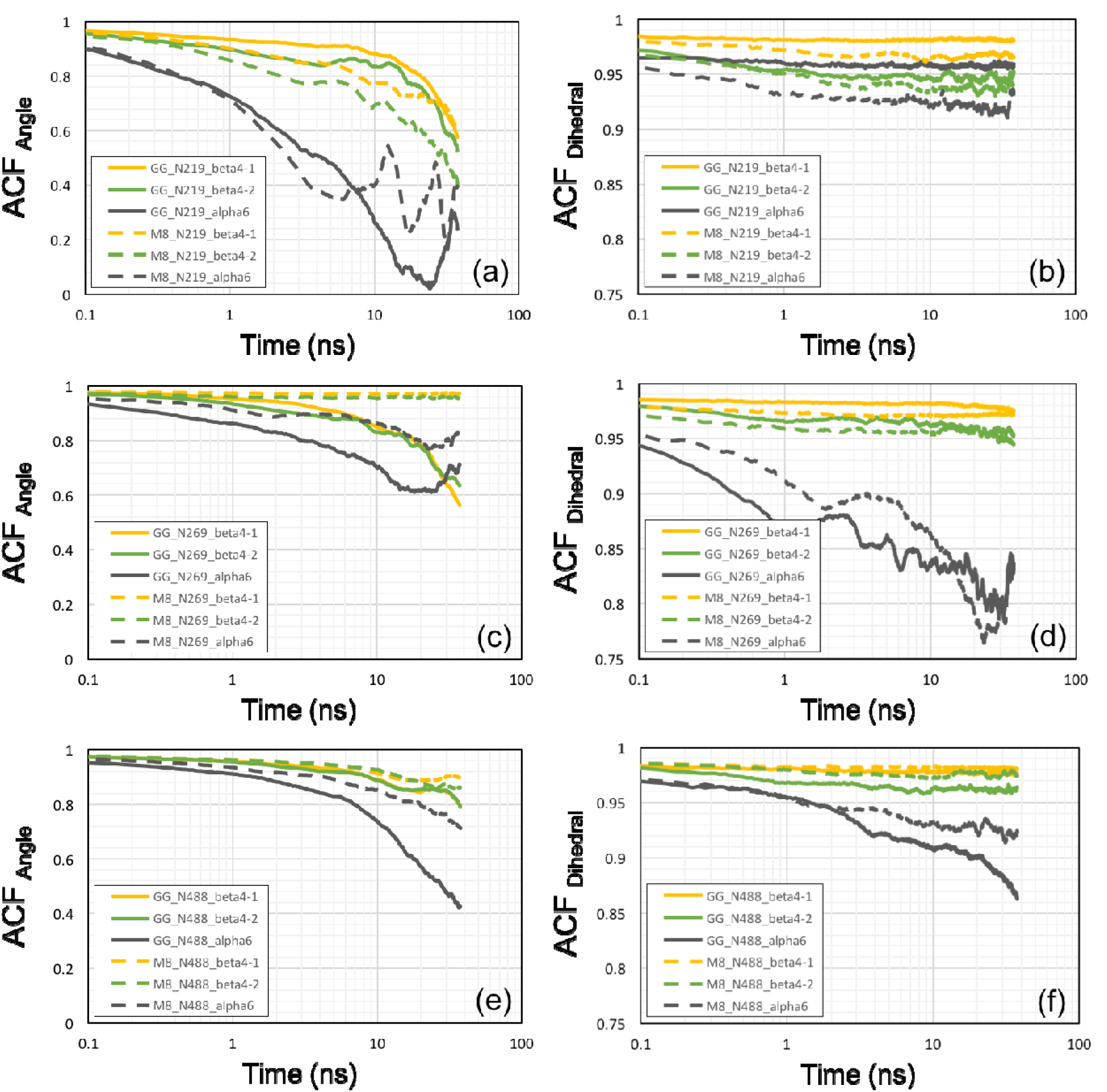
Autocorrelation function analysis of angles and dihedrals at linkage beta4_1, beta4_2, and alpha6 for MAN8 and GnGnXF^3^ at ACE2 fragment glycosylation sites in semi-log lots. Glycans at N219 (a-b) and at N269 (c-d) shows typical behaviors, and glycan at N488 (e-f) are the ones directly interacting with ANaF^6^ on RBD. Dashed lines are the dynamic motions of MAN8, and solid lines are the dynamic motions of GnGnXF^3^.

In addition to hydrogen bonding, we find that electrostatic and Lennard Jones interactions contribute to the binding between ACE2 and RBD. These interactions are plotted in Figure 8 with subplots 8a-d corresponding to the same variants as before. The y-axis corresponds to the interaction energy between the ACE2 and RBD groups with the yellow line corresponding to Coulombic interactions and blue being short range Lennard Jones energies. Interestingly, it appears that at very short distances the electrostatic interaction is more important (more negative interaction potential) than the Lennard Jones interaction; this reverses at intermediate distances (1-2 nm from close contact) where the two lines cross for most of the systems. In some cases, there is a recrossing before the lines essentially merge and the interaction dies out. The glycosylated systems show a similar extension in interaction energies as in the hydrogen bonds, roughly 2-3 nm for the A1Fr^M8^/SpFr and 1 nm for A2Fr^GG^/SpFr. A1 variants demonstrate a differently shaped interaction curve than A2 variants for both glycosylated and aglycosylated systems, this can also be attributable to difference in starting orientation and zinc coordination.

**Figure 8:**
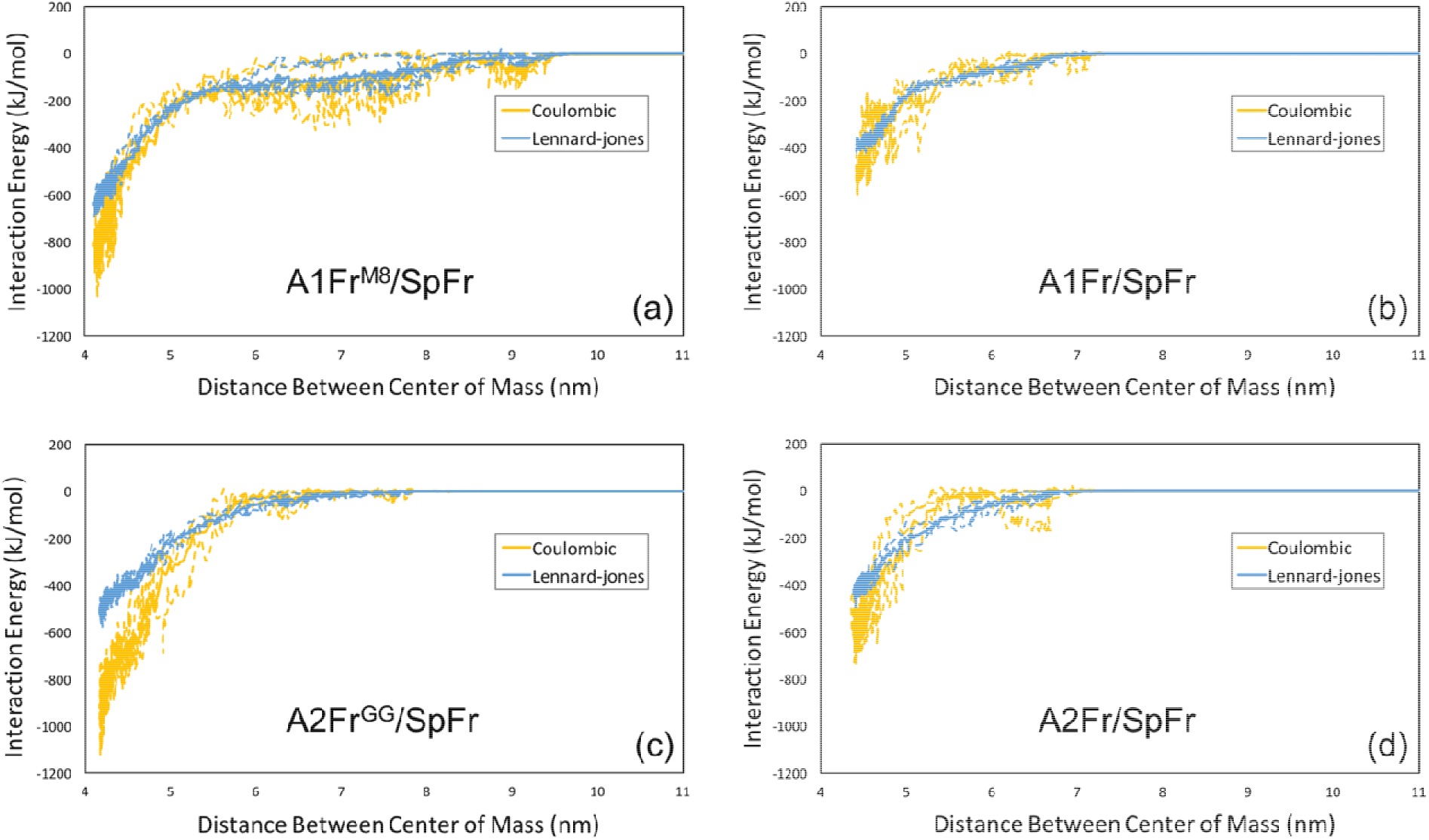
Lennard Jones and electrostatic energies. a) Man8 Glycosylated A1Fr^M8^/SpFr. b) Aglycosylated Structure A1Fr/SpFr. c) GnGnXF Glycosylated A2Fr^GG^/SpFr. d) Aglycosylated Structure A2Fr/SpFr.

Visual inspection of the starting configurations of the two systems shows a difference in RBD alignment in the binding pocket. To evaluate if this difference was due to a rocking motion of the RBD or was caused by differences in the glycans a principal component analysis (PCA) was performed on the trajectories from our previous publication (2) to determine the dominant motions of the RBD. Results of the PCA are presented in Figure 9 and S12-S15. Figures 9a and 9b show still structures with arrows indicating direction of projected motion from the dominant principal component. Corresponding video files are available in SI along with time dependence and pair-wise plots of principal components (Figures S12-S15). Figure 9a shows the motion of the spike fragment from A1Fr^M8^/SpFr is a scissoring between helices and oscillation of the turn at the top of the structure. Figure 9b shows a similar motion, but the oscillation of the turn is missing due to the formation of a helix at that site. This structural change comes from the stable structure after 75 ns simulation due to differences between the glycans and ACE2 interaction. Figures 9c and d show cumulative variance vs number of principal components for A1Fr^M8^/SpFr and A2Fr^GG^/SpFr respectively. This clearly shows that most of the variance is explained by the first principal component (~90% and ~96% for A1Fr^M8^/SpFr and A2Fr^GG^/SpFr, respectively).

**Figure 9:**
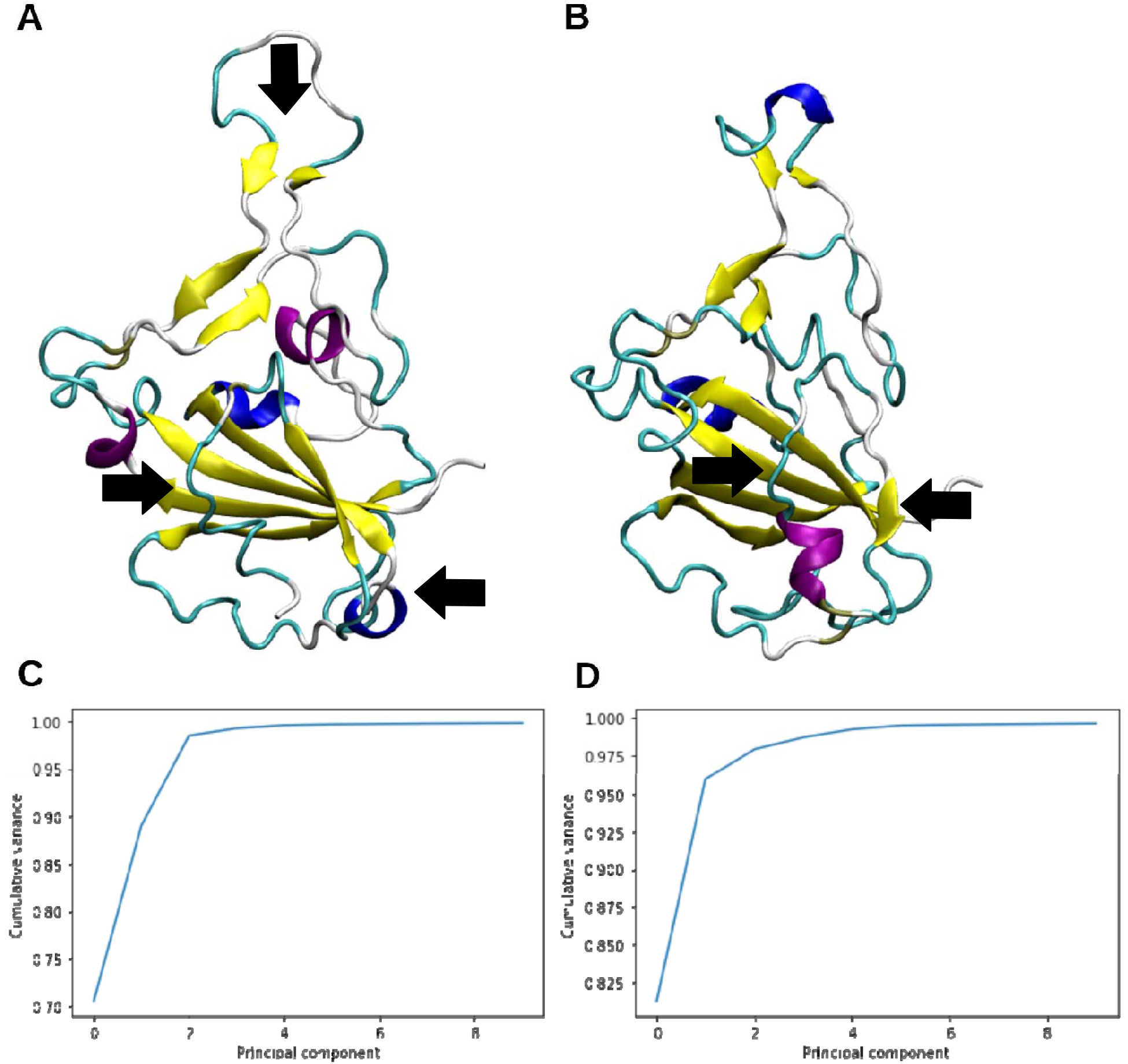
First Principal Component (PC1) projected motion and cumulative variance. A) PC1 projected motion for A1Fr^M8^/SpFr. B) PC1 projected motion for A2Fr^GG^/SpFr C)Principal component cumulative variance A1Fr^M8^/SpFr. D) Principal component cumulative variance A2Fr^GG^/SpFr. Arrows indicate contraction.

To determine whether changes in binding affinity due to deglycosylation can be observed experimentally, we performed biolayer interferometry using ACE2-Fc and RBD with and without removal of N-glycans. Biolayer interferometry is an optical technique that measures biomolecular interactions by detecting changes in the interference pattern of reflected light from a surface before and after binding (57). The response is measured as a shift in wavelength in units of nm. Figure 10a shows that deglycosylation of proteins via PNGase F treatment results in slightly lower bands on an SDS-PAGE gel, as expected from the smaller protein sizes following glycan removal. We then performed biolayer interferometry on ACE2-Fc and RBD that are either both deglycosylated or glycosylated (Figure 10b-d). To do this, ACE2-Fc was immobilized onto a biosensor using the Fc tag and placed in a solution containing the RBD analyte. Steady state analysis was performed on the response using a 1:1 Langmuir binding model, where the response indicates the shift in interference patterns caused by analyte binding (Figure 10d). Glycosylated ACE2-Fc and RBD have a binding affinity, K_D_, of 30 nM, which is similar to values reported by other groups (34,58). Deglycosylation of ACE2-Fc and RBD results in a 2- to 3-fold increase in binding affinity to 77 nM. From the increase in binding affinity, the magnitude of the binding energy decreases by 2.3 kJ/mol following removal of N-linked glycans. This is consistent with our simulation results that predicts that less pulling force is required to break the protein interactions after deglycosylation.

**Figure 10.**
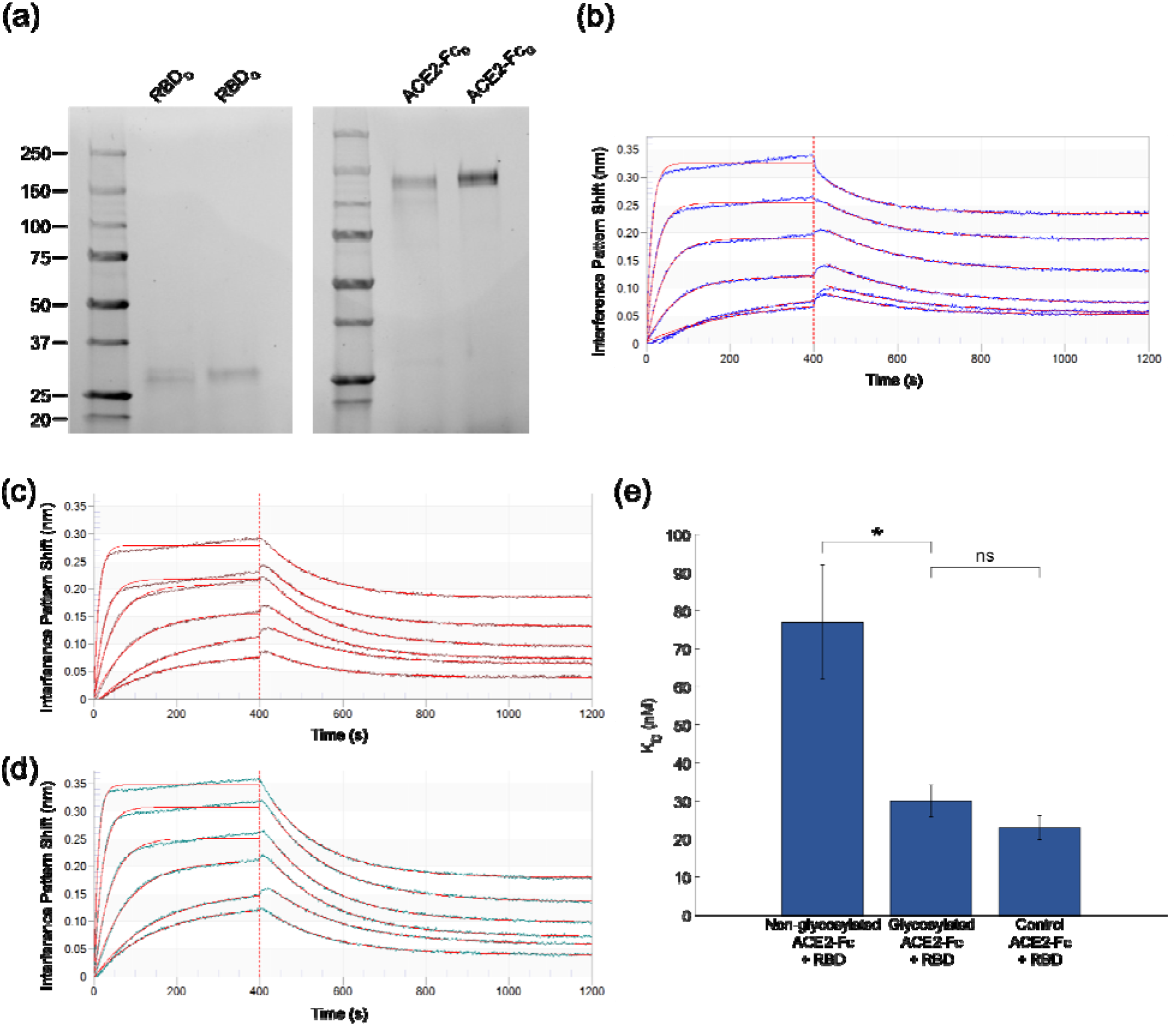
Biolayer Interferometry on glycosylated and deglycosylated ACE2-Fc and RBD. (a) SDS-PAGE on ACE2-Fc and RBD with and without PNGase F treatment. A total of 1 μg of protein is loaded onto each lane. Subscript D indicates deglycosylated proteins, and subscript G indicates glycosylated proteins. (b, c, d) Biolayer interferometry response for (b) deglycosylated ACE2-Fc and RBD, (c) glycosylated ACE2-Fc and RBD, and (d) glycosylated ACE2-Fc and RBD without glyco buffer 2 and incubation at 37°C. Red lines are the fits to the raw data shown in blue, brown, and green, respectively. Error bars represent standard error. * indicates p < 0.05. “ns” indicates not significant (p > 0.05). Probability values were calculated using one-way ANOVA followed by Tukey’s test.

## Discussion

Detailed mechanistic studies of binding interaction events can improve our understanding of how specific changes to proteins affect binding strength. Differences in binding dissociation rate could have implications in infectivity (59–61). Viral protein and host receptor interactions are complex due to the interplay between interaction types, different degrees of motion during a binding event, as well as the role of glycans in shielding or strengthening receptor binding. SARS-CoV-2 spike protein and ACE2 interactions are no different. Understanding the implications of different glycans on the binding behavior of spike could prove useful as more variants emerge with potentially different glycosylation patterns. Recent studies have shown experimentally and computationally that the ACE2 and RBD of coronavirus spike fragments have different binding strengths and dissociation rates when they are glycosylated vs nonglycosylated. (33,34).

Previous computational efforts focused on the binding difference between SARS-CoV-1 and SARS-CoV-2 with glycan interactions modeled by a generic pentasaccharide (32). Their analysis focused on the difference in binding strengths and protein contacts between RBD^CoV1^ and RBD^CoV2^. Our results are in alignment with this trend of stronger interactions caused by the glycans but go further in the analysis of the mechanisms behind this stronger interaction and evaluate more realistic glycan models.

First, our results clearly show that the glycans result in stronger and longer ranged interactions that get extended by a catch-slip mechanism between the glycans, i.e., a hydrogen bond breaks and another one at larger distance takes its place. This catch-slip behavior is clearly seen in the hydrogen bonding maps shown in Figure 4. The catch-slip behavior is a result of the original hydrogen bond interactions that are present relaxing and then reforming later. Analysis of A1Fr^M8^/SpFr in Figure 4a clearly shows the relaxation and reformation of glycan contributed hydrogen bonds. This behavior can be attributed to the increased flexibility of the glycans which increases the ability for these late-stage hydrogen bonds to form due to both increased contacts and increased ability to extend through solution. The different structures of MAN8 and GnGnXF^3^ also contribute to the different hydrogen bond interactions between an ACE2 glycan and RBD glycan. The flatter MAN8 allows more hydrogen bonds between MAN8 and ANaF^6^, therefore causing more glycan-glycan and glycan-protein interactions during pulling for A1Fr^M8^/SpFr than for A2Fr^GG^/SpFr. Angle and dihedrals motions are less flexible for MAN8 than for GnGnXF^3^, especially for the MAN8 and GnGnXF^3^ glycans that directly interact with ANaF^6^, proving that MAN8 is more constrained by the hydrogen bonds between MAN8 and ANaF^6^. The hydrogen bond map of A2Fr^GG^/SpFr in Figure 4c shows that there is a present, but less pronounced, hydrogen bond formation between the glycans. The distance extension is seen clearly in the pull force vs center of mass distances (Figure 3) as well as the interaction energies vs center of mass distances (Figure 8), where the glycosylated structures have their interaction distance extended by as much as 2 nm. This extension can be clearly attributed to the glycans when compared against the hydrogen bond map in Figure 4.

Second, an analysis of hydrogen bond occupancy elucidates that the glycans not only result in secondary binding motifs, but also strengthen and extend the existing protein-protein interactions. This is most clearly seen in the % occupancy numbers for the A2Fr^GG^/SpFr structure, with an increase of several percent in most of the top hydrogen bonds. This trend is also present in A1Fr^M8^/SpFr when looking at the top protein-protein interactions such as RBD-GLY167:ACE2-LYS519 showing an increase of over 3%. This strengthening of the protein-protein hydrogen bonds may be a result of the extra stabilization in the RBD structure provided by the glycan. That the glycans strengthen the interactions is consistent with our biolayer interferometry results. A frequent interaction point of interest is the N-glycosylation site ASN90 on ACE2 and GLN409 and THR415 of the spike RBD. Our results suggest a strong interaction in a nearby site ACE2-TYR249 (equivalent to TYR83 in standard numbering) and RBD-ASN152 (equivalent to ASN 487) for all variants studied. This interaction agrees with previous results suggesting a long interaction at this site due to the flexibility of the RBD loop (32). It is interesting to note that this interaction is seemingly not affected by the glycan as it pertains to % occupancy.

It is necessary to comment on the difference in starting orientation of the RBD and the ACE2 between the two different starting truncations. By taking the final structure of the simulations from our previous study, it was possible that this resulted in a lower probability starting orientation. A principal component analysis was performed (Fig 10) to verify that the starting structures were truly the dominant orientation from our previous paper and not just an unlucky snapshot of a less favorable state. These results show that the dominant motion from the highest principal component is scissoring of helices and oscillation of a turn and not the rocking of the spike fragment. This suggests that the structure was stable in the ACE2 binding pocket and that the difference in starting structure is due to the differences between glycosylation and the effect of Zn^2+^on the stability of ACE2. Figures 10 a, b clearly show the structural changes resulting from these interactions. These structural changes result in differences in the interaction behavior as seen by a slight 1nm extension of interaction energies as shown in Fig. 10 b,d.

## Conclusion

We have expanded on our previously developed model of fully glycosylated ACE2-Fc and SARS-CoV-2 spike protein fragments through the investigation of the binding strength and role of glycosylation on binding between these groups. This investigation provides further evidence that the binding between SARS-CoV-2 spike and ACE2 receptor are aided by the glycosylation on each protein. We found that for multiple complex glycan types the interactions between RBD and ACE2 were strengthened and longer ranged. Protein-protein interactions were extended due to the increased stability provided by the glycans and binding strength is affected by a catch-slip behavior between the glycans. These computational results were corroborated by experimental evidence that the magnitude of the binding energy is decreased for deglycosylated proteins. Further work in analyzing the larger fragments of spike will be necessary for a more realistic model of RBD stability in order to address effects of mutations.

