## Supplemental Data for "SARS-Cov-2 Spike binding to ACE2 is stronger and longer ranged due to glycan interaction"

X equal first authors

### 1. Amino Acid Sequences

**Variant 1: ACE2WildType(18-740)-SSERKCCVE-IgG1Fc(109-330)-SEKDEL**

NCBI Reference Sequence ID: NP_001358344.1 (ACE2); UniProtKB Sequence ID: P01857 (IgG1Fc_human corresponds to amino acid residues 109-330)

PDB codes: 6M17 or 6M18 (ACE2), 1R42 (Zn + coordinating residues + coordinating water), 3SGJ (Fc)

```
  1 - QSTIEEQAKTFLDKFNHEAEDLFYQSSLASWNYNTNITEENVQNMNNAGD -  50
 51 - KWSAFLKEQSTLAQMYPLQEIQNLTVKLQLQALQQNGSSVLSEDKSKRLN - 100
101 - TILNTMSTIYSTGKVCNPDNPQECLLLEPGLNEIMANSLDYNERLWAWES - 150
151 - WRSEVGKQLRPLYEEYVVLKNEMARANHYEDYGDYWRGDYEVNGVDGYDY - 200
201 - SRGQLIEDVEHTFEEIKPLYEHLHAYVRAKLMNAYPSYISPIGCLPAHLL - 250
251 - GDMWGRFWTNLYSLTVPFGQKPNIDVTDAMVDQAWDAQRIFKEAEKFFVS - 300
301 - VGLPNMTQGFWENSMLTDPGNVQKAVCHPTAWDLGKGDFRILMCTKVTMD - 350
351 - DFLTAHHEMGHIQYDMAYAAQPFLLRNGANEGFHEAVGEIMSLSAATPKH - 400
401 - LKSIGLLSPDFQEDNETEINFLLKQALTIVGTLPFTYMLEKWRWMVFKGE - 450
451 - IPKDQWMKKWWEMKREIVGVVEPVPHDETYCDPASLFHVSNDYSFIRYYT - 500
501 - RTLYQFQFQEALCQAAKHEGPLHKCDISNSTEAGQKLFNMLRLGKSEPWT - 550
551 - LALENVVGAKNMNVRPLLNYFEPLFTWLKDQNKNSFVGWSTDWSPYADQS - 600
601 - IKVRISLKSALGDKAYEWNDNEMYLFRSSVAYAMRQYFLKVKNQMILFGE - 650
651 - EDVRVANLKPRISFNFFVTAPKNVSDIIPRTEVEKAIRMSRSRINDAFRL - 700
701 - NDNSLEFLGIQPTLGPPNQPPVSSSERKCCVECPPCPAPELLGGPSVFLF - 750
751 - PPKPKDTLMISRTPEVTCVVVDVSHEDPEVKFNWYVDGVEVHNAKTKPRE - 800
801 - EQYNSTYRVVSVLTVLHQDWLNGKEYKCKVSNKALPAPIEKTISKAKGQP - 850
851 - REPQVYTLPPSRDELTKNQVSLTCLVKGFYPSDIAVEWESNGQPENNYKT - 900
901 - TPPVLDSDGSFFLYSKLTVDKSRWQQGNVFSCSVMHEALHNHYTQKSLSL - 950
951 - SPGKSEKDEL
```

Sequence Seq1. ACE2-Fc Variant A1 sequence. ACE2 in orange, linker in red, Fc in grey. Glycosylation sites highlighted in green. Coordinating Zn2+ residues highlighted in pink. Last amino acid kept in truncation for pulling simulation indicated by red highlight.

**Variant 2: ACE2Mutant(18-740,H374N,H378N)-SSERKCCVE-IgG1Fc(109-330)**

NCBI Reference Sequence ID: NP_001358344.1 (ACE2), UniProtKB Sequence ID: P01857 (IgG1Fc_human corresponds to amino acid residues 109 – 330)

PDB codes: 6M17 or 6M18 (ACE2), 3SGJ (Fc)

```
  1 - QSTIEEQAKTFLDKFNHEAEDLFYQSSLASWNYNTNITEENVQNMNNAGD -  50
 51 - KWSAFLKEQSTLAQMYPLQEIQNLTVKLQLQALQQNGSSVLSEDKSKRLN - 100
101 - TILNTMSTIYSTGKVCNPDNPQECLLLEPGLNEIMANSLDYNERLWAWES - 150
151 - WRSEVGKQLRPLYEEYVVLKNEMARANHYEDYGDYWRGDYEVNGVDGYDY - 200
201 - SRGQLIEDVEHTFEEIKPLYEHLHAYVRAKLMNAYPSYISPIGCLPAHLL - 250
251 - GDMWGRFWTNLYSLTVPFGQKPNIDVTDAMVDQAWDAQRIFKEAEKFFVS - 300
301 - VGLPNMTQGFWENSMLTDPGNVQKAVCHPTAWDLGKGDFRILMCTKVTMD - 350
351 - DFLTAHNEMGNIQYDMAYAAQPFLLRNGANEGFHEAVGEIMSLSAATPKH - 400
401 - LKSIGLLSPDFQEDNETEINFLLKQALTIVGTLPFTYMLEKWRWMVFKGE - 450
451 - IPKDQWMKKWWEMKREIVGVVEPVPHDETYCDPASLFHVSNDYSFIRYYT - 500
501 - RTLYQFQFQEALCQAAKHEGPLHKCDISNSTEAGQKLFNMLRLGKSEPWT - 550
551 - LALENVVGAKNMNVRPLLNYFEPLFTWLKDQNKNSFVGWSTDWSPYADQS - 600
601 - IKVRISLKSALGDKAYEWNDNEMYLFRSSVAYAMRQYFLKVKNQMILFGE - 650
651 - EDVRVANLKPRISFNFFVTAPKNVSDIIPRTEVEKAIRMSRSRINDAFRL - 700
701 - NDNSLEFLGIQPTLGPPNQPPVSSSERKCCVECPPCPAPELLGGPSVFLF - 750
751 - PPKPKDTLMISRTPEVTCVVVDVSHEDPEVKFNWYVDGVEVHNAKTKPRE - 800
801 - EQYNSTYRVVSVLTVLHQDWLNGKEYKCKVSNKALPAPIEKTISKAKGQP - 850
851 - REPQVYTLPPSRDELTKNQVSLTCLVKGFYPSDIAVEWESNGQPENNYKT - 900
901 - TPPVLDSDGSFFLYSKLTVDKSRWQQGNVFSCSVMHEALHNHYTQKSLSL - 950
951 - SPGK
```

Sequence Seq2. ACE2-Fc variant 2 sequence. ACE2 in orange, linker in red, Fc in grey. Glycosylation sites highlighted in green. Mutated residues highlighted in pink. Last amino acid kept in truncation for pulling simulation indicated by red highlight.

**SpFr (crystallized residues only):**

NCBI Reference Sequence ID: YP_009724390.1(SpFr corresponds to amino acid residues 336 – 518)
PDB codes: 6M17

```
  1 - CPFGEVFNATRFASVYAWNRKRISNCVADYSVLYNSASFSTFKCYGVSPT -  50
 51 - KLNDLCFTNVYADSFVIRGDEVRQIAPGQTGKIADYNYKLPDDFTGCVIA - 100
101 - WNSNNLDSKVGGNYNYLYRLFRKSNLKPFERDISTEIYQAGSTPCNGVEG - 150
151 - FNCYFPLQSYGFQPTNGVGYQPYRVVVLSFELL
```

Sequence Seq3. Spike fragment (SpFr) sequence. Glycosylation site highlighted in green.

#### 2. Molecular Dynamics Pulling

##### 2.1 Pull Force vs Pull distance

As described in the manuscript, we analyzed pulling speeds of 10 nm/ns, 5nm/ns, and 1 nm/ns. Here we present the pull force vs pull distance plots for these three different pull rates. As expected pull force increases with pulling rate (blue, orange, green), and the peak force is clearly higher in the glycosylated states by roughly 250, 500, and 600 kJ /mol /nm.

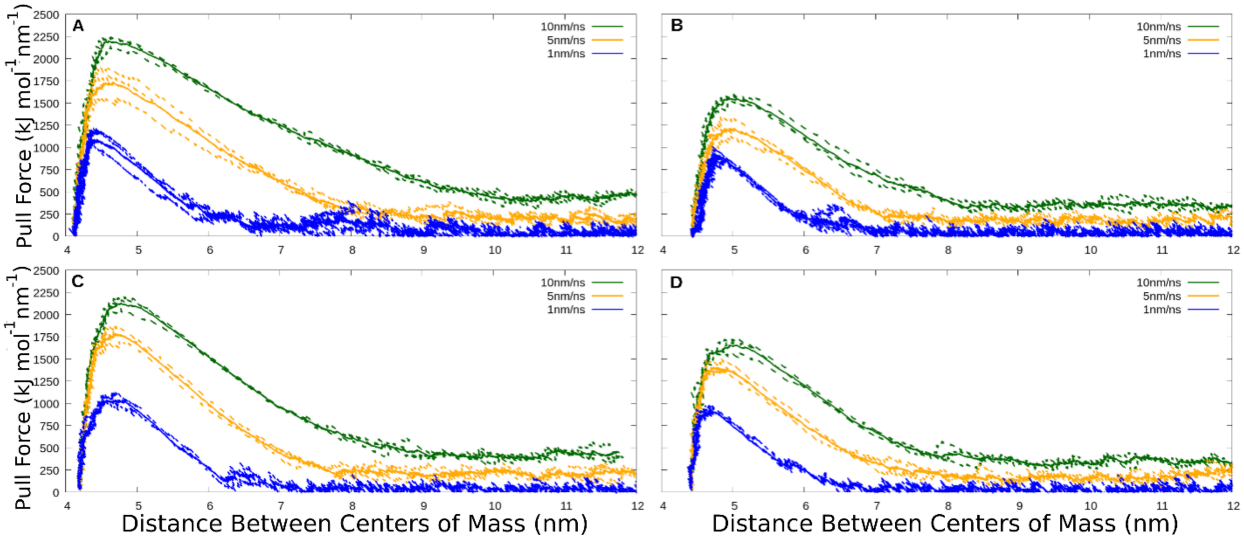

Figure S1: Traces of pull force versus distance. A) Man8 Glycosylated A1Fr$^{M8}$/SpFr. B) Non-glycosylated Structure A1Fr/SpFr. C) GnGnXF Glycosylated A2Fr$^{GG}$/SpFr. D) Non-glycosylated Structure A2Fr/SpFr. Different colors indicate different pull rate. Dashed lines are individual replicas, solid lines are averages.

### 3. Hydrogen Bonding

#### 3.1 Hydrogen Bonding Script

Hydrogen bonding maps are generated from a python script using external packages, numpy, pandas, matplotlib, gromacs and seaborn. The gromacs python package is used to load the .xpm bitmap generated from the gromacs hbond command and save it as a python array. The log and index files from the gromacs hbond command are then sorted and used to generate labels for the previously generated array. For plotting simplicity, the corresponding array and labels are converted to a pandas data frame and plotted using the seaborn heatmap. The transformations made before plotting include a % occupancy calculation attained by calculating the number of 1s in the array divided by the number of columns in the row and multiplied by 100, and some conditional dataframe rearrangement based on interaction type. This rearrangement was used to generate different colors for each interaction type by either multiplying values by -1 or 2 depending on the interaction involved.

#### 3.2 Hydrogen Bonding Maps

As described in the manuscript we analyzed the hydrogen bonding interactions between ACE2 and Spike RBD proteins. Interactions were calculated and mapped as described above. Manuscript figure 4 was shown without labels due to lack of available resolution. Full scale images with labels are shown below in Figure S1-S4. Data is from 1 nm/ns pull rate for each system. Colors indicate interaction type. White: protein-protein, yellow: protein-glycan, and magenta: glycan-glycan.

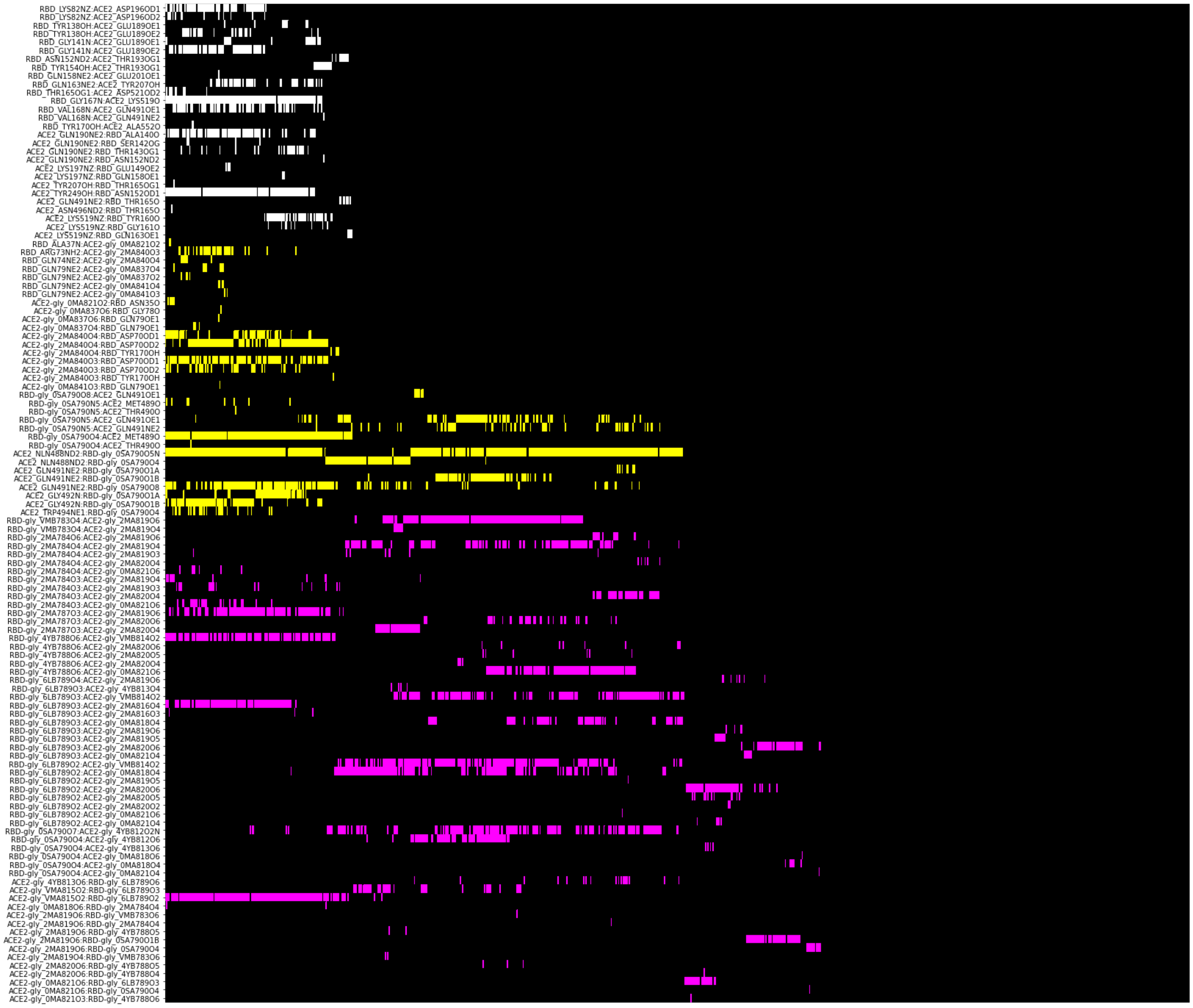

**Figure S2.** Hydrogen bonding donor:acceptor pairs vs simulated time for A1Fr$^{M8}$/SpFr. 1 ns / nm pulling speed. Colors indicate interaction type: White: protein-protein, Yellow: protein-glycan, Magenta: glycan-glycan

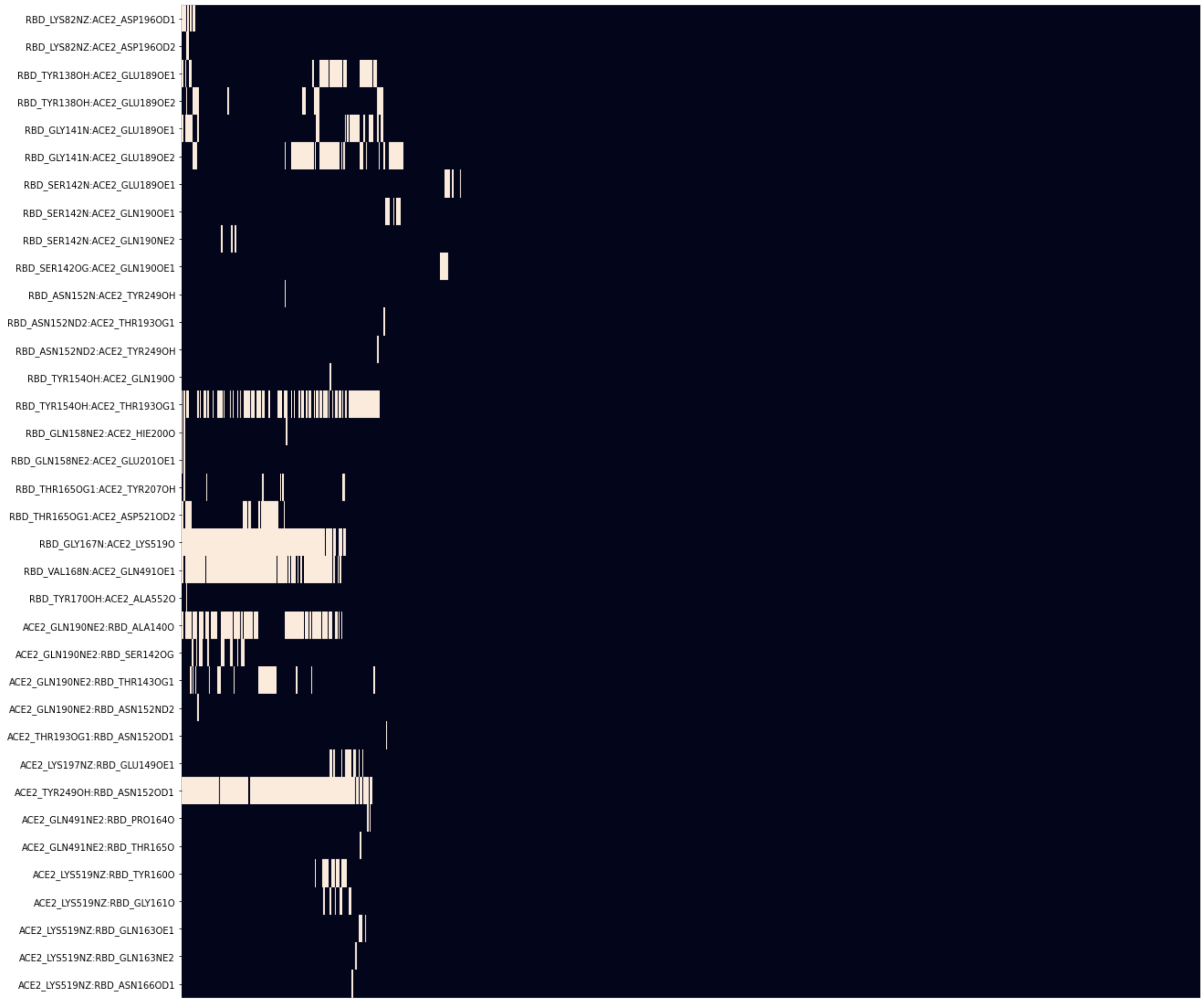

**Figure S3.** Hydrogen bonding donor:acceptor pairs vs simulated time for A1Fr/SpFr. 1 ns / nm pulling speed. Colors indicate interaction type: White: protein-protein

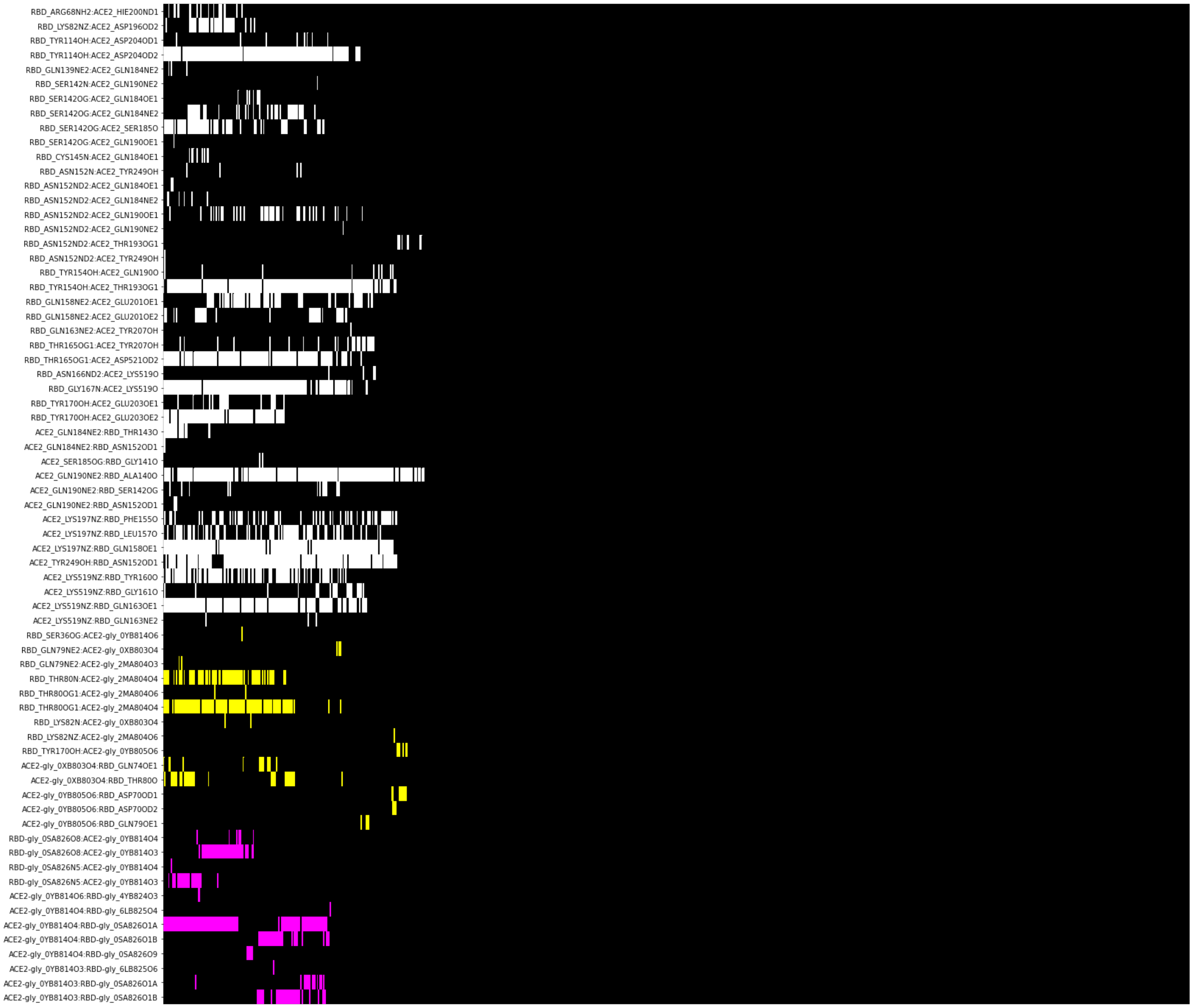

**Figure S4.** Hydrogen bonding donor:acceptor pairs vs simulated time for A2Fr$^{GG}$/SpFr. 1 ns / nm pulling speed. Colors indicate interaction type: White: protein-protein, Yellow: protein-glycan, Magenta: glycan-glycan

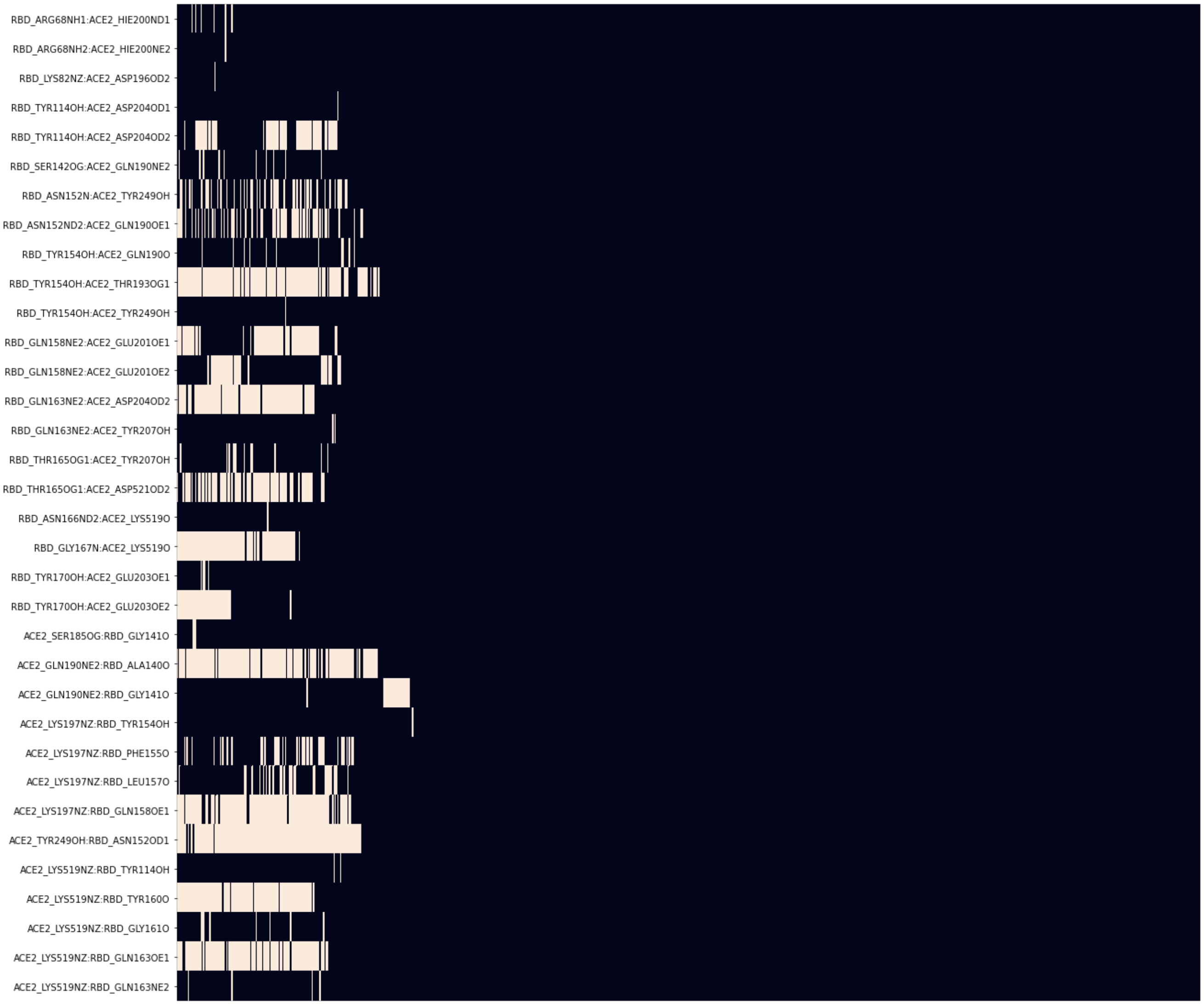

**Figure S5.** Hydrogen bonding donor:acceptor pairs vs simulated time for A2Fr/SpFr. 1 ns / nm pulling speed. Colors indicate interaction type: White: protein-protein

#### 3.3 Hydrogen Bond Occupancy

As described in the manuscript, the top 25 hydrogen bonds by occupancy were listed for all four system, and the top 5 donor (blue):acceptor (pink) pairs by occupancy were highlighted in the four configurations where RBD with and without ANaF[6] started to be pulled away from the ACE2 fragment. Figure S5-S8 correspond to $A1Fr^{M8}$/SpFr, A1Fr/SpFr, $A2Fr^{GG}$/SpFr, A2Fr/SpFr respectively.

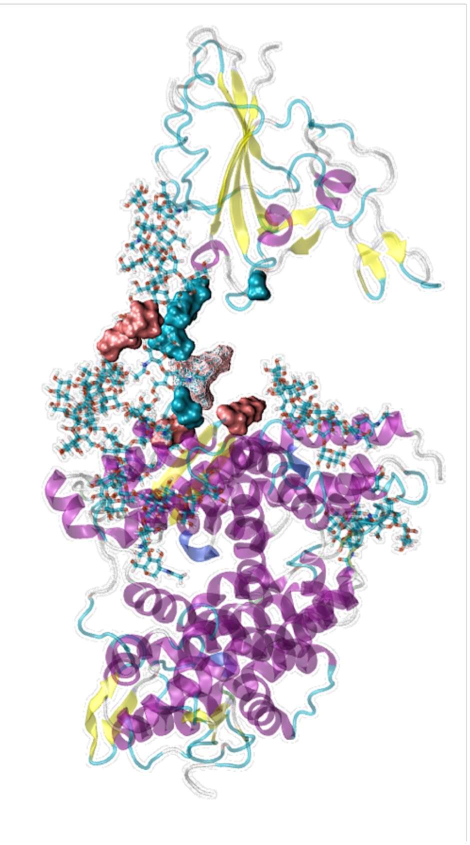

**Glycosylated ACE2_Man8 and RBD Top 25 Hbonds**

| Donor Acceptor Pair (Donor : Acceptor) | Total Occupancy (%) |
|---|---|
| RBD-gly_6LB789O2:ACE2-gly_VMA815O2 | 24.6 |
| ACE2_NLN488ND2:RBD-gly_0SA790O5N | 20.2 |
| RBD-gly_0SA790O4:ACE2_MET489O | 19.8 |
| RBD_GLY167N:ACE2_LYS519O | 18.3 |
| RBD-gly_4YB788O6:ACE2-gly_VMB814O2 | 18.1 |
| RBD-gly_6LB789O3:ACE2_gly_2MA816O4 | 17.9 |
| ACE2_TYR249OH:RBD_ASN152OD1 | 17.9 |
| ACE2-gly_2MA840O4:RBD_ASP70OD1 | 15.6 |
| RBD-gly_2MA787O3:ACE2-gly_2MA819O6 | 15.4 |
| ACE2_GLN190NE2:RBD_ALA140O | 14.2 |
| ACE2_GLY492N:RBD-gly_0SA790O1A | 13.8 |
| ACE2_GLY491NE2:RBD-gly_0SA790O8 | 13.0 |
| RBD_VAL168N:ACE2_GLN491OE1 | 11.3 |
| RBD-gly_2MA784O3:ACE2-gly_2MA819O3 | 11.0 |
| ACE2-gly_2MA840O3:RBD_ASP70OD1 | 11.0 |
| ACE2_TRP494NE1:RBD-gly_0SA790O4 | 8.7 |
| RBD_GLY141N:ACE2_GLU189OE1 | 8.1 |
| ACE2-gly_VMB814O2:RBD-gly_6LB789O2 | 7.3 |
| ACE2_GLY492N:RBD-gly_0SA790O1B | 7.2 |
| RBD_TYP138OH:ACE2_GLU189OE2 | 6.3 |
| ACE2-gly_2MA840O3:RBD_ASP70OD2 | 5.4 |
| RBD_GLY141N:ACE2_GLU189OE2 | 4.7 |
| RBD_LYS82NZ:ACE2_ASP196OD1 | 4.3 |
| ACE2-gly_2MA819O3:RBD-gly_6LB789O2 | 4.3 |
| RBD_TYR154OH:ACE2_THR193OG1 | 4.0 |

**Figure S6.** Hydrogen bond donor:acceptor pairs and occupancy for Man8 glycosylated A1Fr$^{M8}$/SpFr. Table colors indicate interaction type: White: protein-protein, Yellow: protein-glycan, Magenta: glycan-glycan.

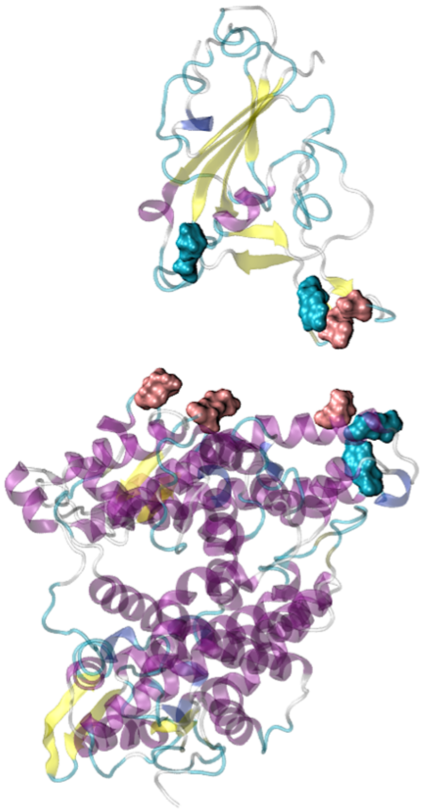

| Aglycosylated ACE2_MAN8 and RBD Top 25 Hbonds | |
|---|---|
| **Donor Acceptor Pair (Donor : Acceptor)** | **Total Occupancy (%)** |
| ACE2_TYR249OH:RBD_ASN152OD1 | 18.0 |
| RBD_GLY167N:ACE2_LYS519O | 15.6 |
| RBD_VAL168N:ACE2_GLN491OE1 | 14.2 |
| RBD_TYR154OH:ACE2_THR193OG1 | 11.2 |
| ACE2_GLN190NE2:RBD_ALA140O | 10.5 |
| RBD_GLY141N:ACE2_GLU189OE2 | 7.4 |
| RBD_TYR138OH:ACE2_GLU189OE1 | 4.8 |
| RBD_GLY141N:ACE2_GLU189OE1 | 3.7 |
| RBD_THR165OG1:ACE2_ASP521OD2 | 3.6 |
| ACE2_GLN190NE2:RBD_THR143OG1 | 3.1 |
| RBD_TYR138OH:ACE2_GLU189OE2 | 2.5 |
| ACE2_LYS519NZ:RBD_TYR160O | 2.1 |
| ACE2_GLN190NE2:RBD_SER142OG | 2.0 |
| ACE2_LYS197NZ:RBD_GLU149OE1 | 1.6 |
| RBD_SER142N:ACE2_GLN190OE1 | 1.1 |
| RBD_LYS82NZ:ACE2_ASP196OD1 | 1.0 |
| RBD_SER142OG:ACE2_GLN190OE1 | 0.9 |
| RBD_THR165OG1:ACE2_TYR207OH | 0.9 |
| ACE2_LYS519NZ:RBD_GLY161O | 0.9 |
| RBD_SER142N:ACE2_GLU189OE1 | 0.7 |
| ACE2_LYS519NZ:RBD_GLN163OE1 | 0.5 |
| RBD_SER142N:ACE2_GLN190NE2 | 0.4 |
| RBD_GLN158NE2:ACE2_HIE200O | 0.4 |
| RBD_LYS82NZ:ACE2_ASP196OD2 | 0.2 |
| RBD_GLN158NE2:ACE2_GLU201OE1 | 0.2 |

**Figure S7.** Hydrogen bond donor:acceptor pairs and occupancy for non-glycosylated structure A1Fr/SpFr. Table colors indicate interaction type: White: protein-protein, Yellow: protein-glycan, Magenta: glycan-glycan.

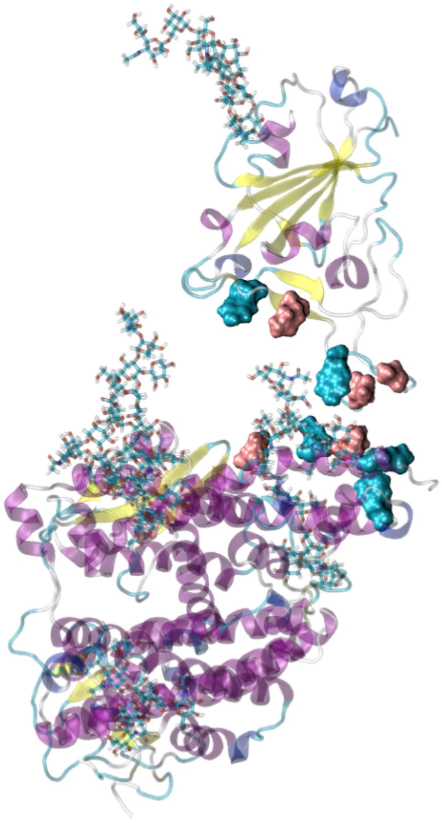

**Glycosylated ACE2_GnGnXF and RBD Top 25 Hbonds**

| Donor Acceptor Pair (Donor : Acceptor) | Total Occupancy (%) |
|---|---|
| ACE2_GLN190NE2:RBD_ALA140O | 23.1 |
| ACE2_LYS197NZ:RBD_GLN158OE1 | 21.6 |
| RBD_TYR154OH:ACE2_THR193OG1 | 21.2 |
| ACE2_TYR249OH:RBD_ASN152OD1 | 20.2 |
| RBD_TYR114OH:ACE2_ASP204OD2 | 18.2 |
| RBD_GLY167N:ACE2_LYS519O | 17.4 |
| ACE2_LYS519NZ:RBD_GLN163OE1 | 17.2 |
| RBD_THR165OG1:ACE2_ASP521OD2 | 16.2 |
| ACE2-gly_0YB814O4:RBD-gly_0SA826O1A | 11.9 |
| RBD_THR80OG1:ACE2-gly_2MA804O4 | 11.7 |
| RBD_TYR170OH:ACE2_GLU203OE2 | 11.0 |
| ACE2_LYS519NZ:RBD_TYR160O | 11.0 |
| ACE2_LYS197NZ:RBD_LEU157O | 8.6 |
| RBD_THR80N:ACE2-gly_2MA804O4 | 8.0 |
| ACE2_LYS197NZ:RBD_PHE155O | 7.9 |
| RBD_GLN158NE2:ACE2_GLU201OE1 | 7.7 |
| RBD_SER142OG:ACE2_SER185O | 7.6 |
| RBD_SER142OG:ACE2_GLN184NE2 | 5.0 |
| RBD-gly_0SA826O8:ACE2-gly_0YB814O3 | 4.9 |
| RBD_ASN152ND2:ACE2_GLN190OE1 | 4.7 |
| RBD_LYS82NZ:ACE2_ASP196OD2 | 4.4 |
| RBD_GLN158NE2:ACE2_GLU201OE2 | 4.1 |
| ACE2-gly_0YB814O3:RBD-gly_0SA826O1B | 4.0 |
| ACE2-gly_0XB803O4:RBD_THR80O | 3.9 |
| ACE2-gly_0YB814O4:RBD-gly_0SA826O1B | 3.6 |

**Figure S8.** Hydrogen bond donor:acceptor pairs and occupancy for GnGnXF[3] glycosylated A2Fr$^{GG}$/SpFr. Table colors indicate interaction type: White: protein-protein, Yellow: protein-glycan, Magenta: glycan-glycan.

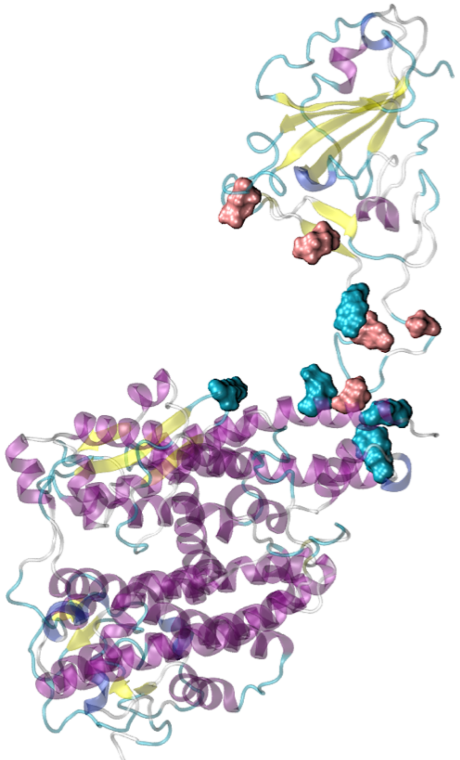

**Aglycosylated ACE2_GnGnXF and RBD Top 25 Hbonds**

| Donor Acceptor Pair (Donor : Acceptor) | Total Occupancy (%) |
|---|---|
| ACE2_TYR249OH:RBD_ASN152OD1 | 19.8 |
| ACE2_GLN190NE2:RBD_ALA140O | 19.2 |
| RBD_TYR154OH:ACE2_THR193OG1 | 18.7 |
| ACE2_LYS197NZ:RBD_GLN158OE1 | 16.8 |
| ACE2_LYS519NZ:RBD_GLN163OE1 | 14.6 |
| ACE2_LYS519NZ:RBD_TYR160O | 14.5 |
| RBD_GLN163NE2:ACE2_ASP204OD2 | 14.0 |
| RBD_GLY167N:ACE2_LYS519O | 12.5 |
| RBD_THR165OG1:ACE2_ASP521OD2 | 11.2 |
| RBD_GLN158NE2:ACE2_GLU201OE1 | 9.7 |
| RBD_TYR114OH:ACE2_ASP204OD2 | 8.6 |
| RBD_ASN152ND2:ACE2_GLN190OE1 | 8.6 |
| RBD_ASN152N:ACE2_TYR249OH | 6.7 |
| RBD_TYR170OH:ACE2_GLU203OE2 | 6.2 |
| ACE2_LYS197NZ:RBD_PHE155O | 5.6 |
| RBD_GLN158NE2:ACE2_GLU201OE2 | 5.2 |
| ACE2_LYS197NZ:RBD_LEU157O | 4.4 |
| ACE2_GLN190NE2:RBD_GLY141O | 3.1 |
| RBD_THR165OG1:ACE2_TYR207OH | 1.7 |
| RBD_TYR154OH:ACE2_GLN190O | 1.5 |
| RBD_SER142OG:ACE2_GLN190NE2 | 1.4 |
| ACE2_LYS519NZ:RBD_GLY161O | 1.1 |
| RBD_ARG68NH1:ACE2_HIE200ND1 | 0.8 |
| RBD_TYR170OH:ACE2_GLU203OE1 | 0.6 |
| ACE2_LYS519NZ:RBD_GLN163NE2 | 0.6 |

**Figure S9.** Hydrogen bond donor:acceptor pairs and occupancy for non-glycosylated structure A2Fr/SpFr. Table colors indicate interaction type: White: protein-protein, Yellow: protein-glycan, Magenta: glycan-glycan.

### 4. Autocorrelation Functions

#### 4.1 angle correlation functions and dihedral correlation functions

As discussed in the manuscript, both glycosylated systems A1Fr$^{M8}$/SpFr and A2Fr$^{GG}$/SpFr have 6 glycosylation sites on the ACE2 fragment: N219, N256, N269, N488, N598, N712. Angle autocorrelation functions (ACF) and dihedral autocorrelation functions were calculated at glycan linkages beta4_1, beta4_2, and alpha6 at all 6 glycosylation sites for both systems. Figure S9 shows all the angle ACF semi-log plots, and Figure S10 shows all the dihedral ACF semi-log plots.

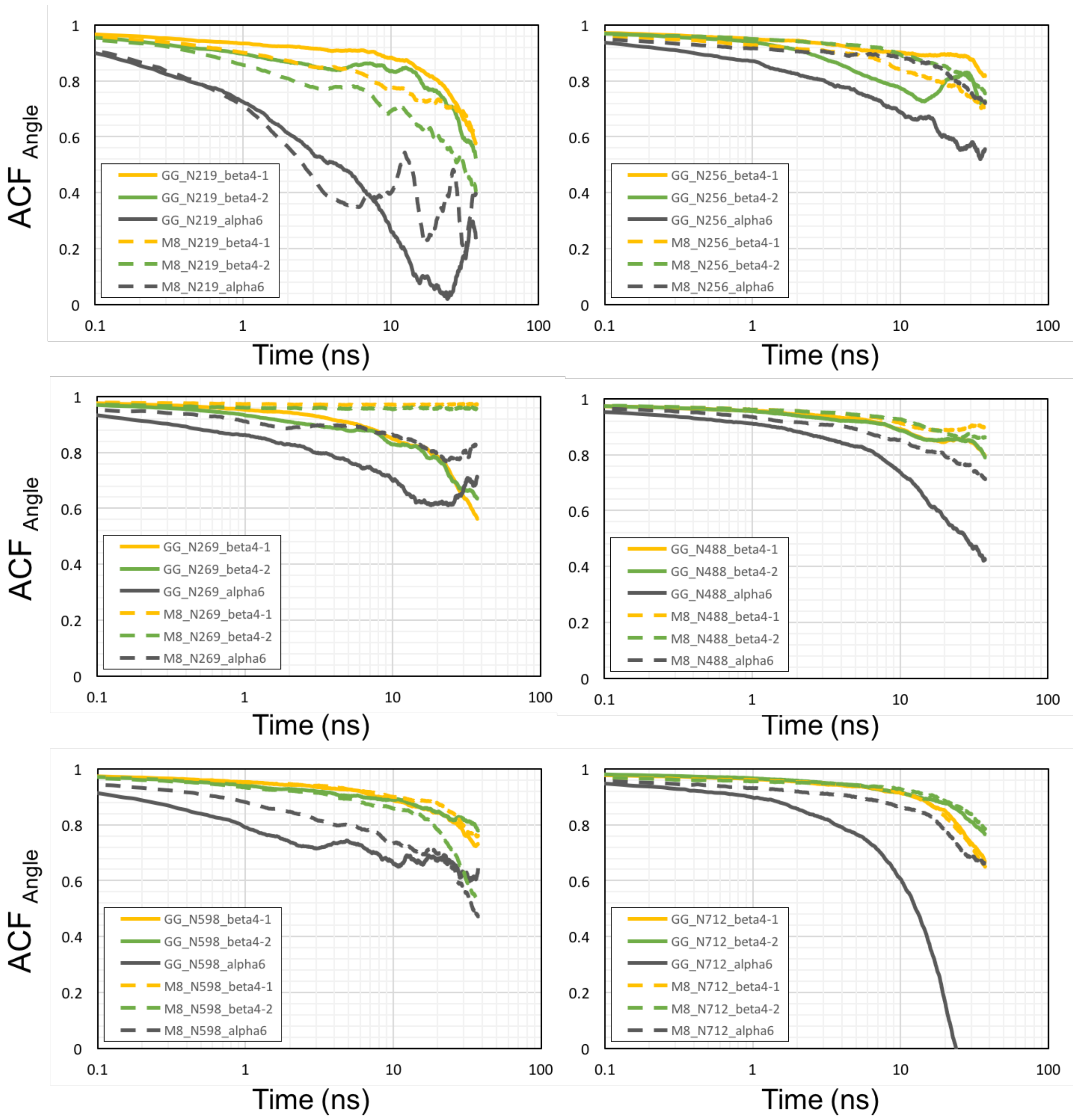

**Figure S10.** Autocorrelation function analysis of angles at linkage beta4_1, beta4_2, and alpha6 of MAN8 and GnGnXF[3] at ACE2 fragment glycosylation sites in semi-log plots. Dashed lines are the dynamic motions of MAN8, and solid lines are the dynamic motions of GnGnXF[3].

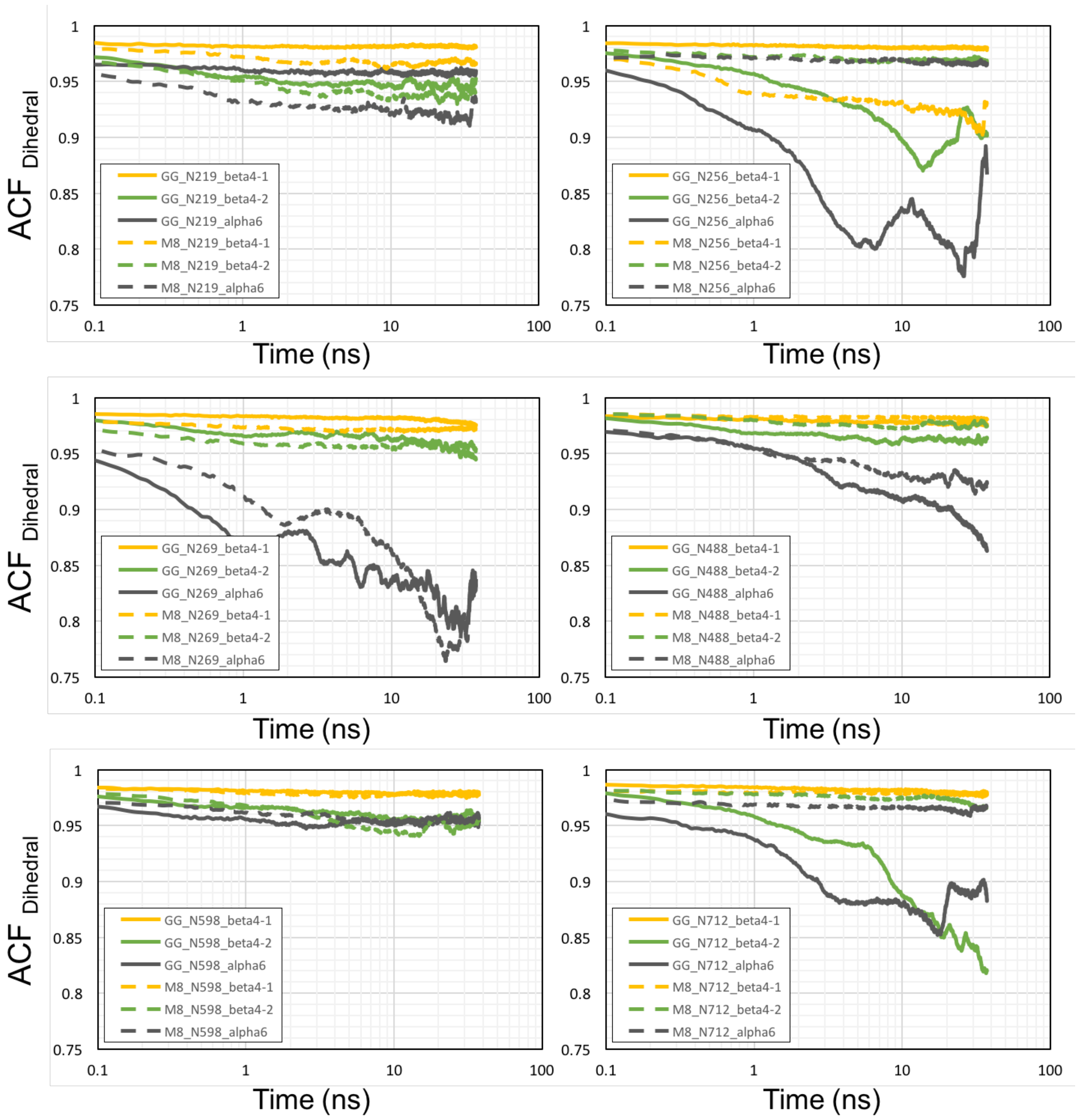

**Figure S11.** Autocorrelation function analysis of dihedrals at linkage beta4_1, beta4_2, and alpha6 of MAN8 and GnGnXF$^3$ at ACE2 fragment glycosylation sites in semi-log plots. Dashed lines are the dynamic motions of MAN8, and solid lines are the dynamic motions of GnGnXF$^3$.

### 5. Principal Component Analysis

#### 5.1 Principal components

As described in the manuscript, PCA was performed on the trajectories from our previous publication to determine the dominant motion of the RBD. The results in the manuscript show that roughly 90% and 95% of the variance in the motion was explained by the first component for the A1 and A2 variants respectively. Here we present the first 5 components, responsible for over 99% of the variance in both systems as a pair-wise interaction plot in Figure S11 and S12 for A1 and A2 respectively, and as PC vs time in figure S13 and S14. Simulation videos are included as supplementary files.

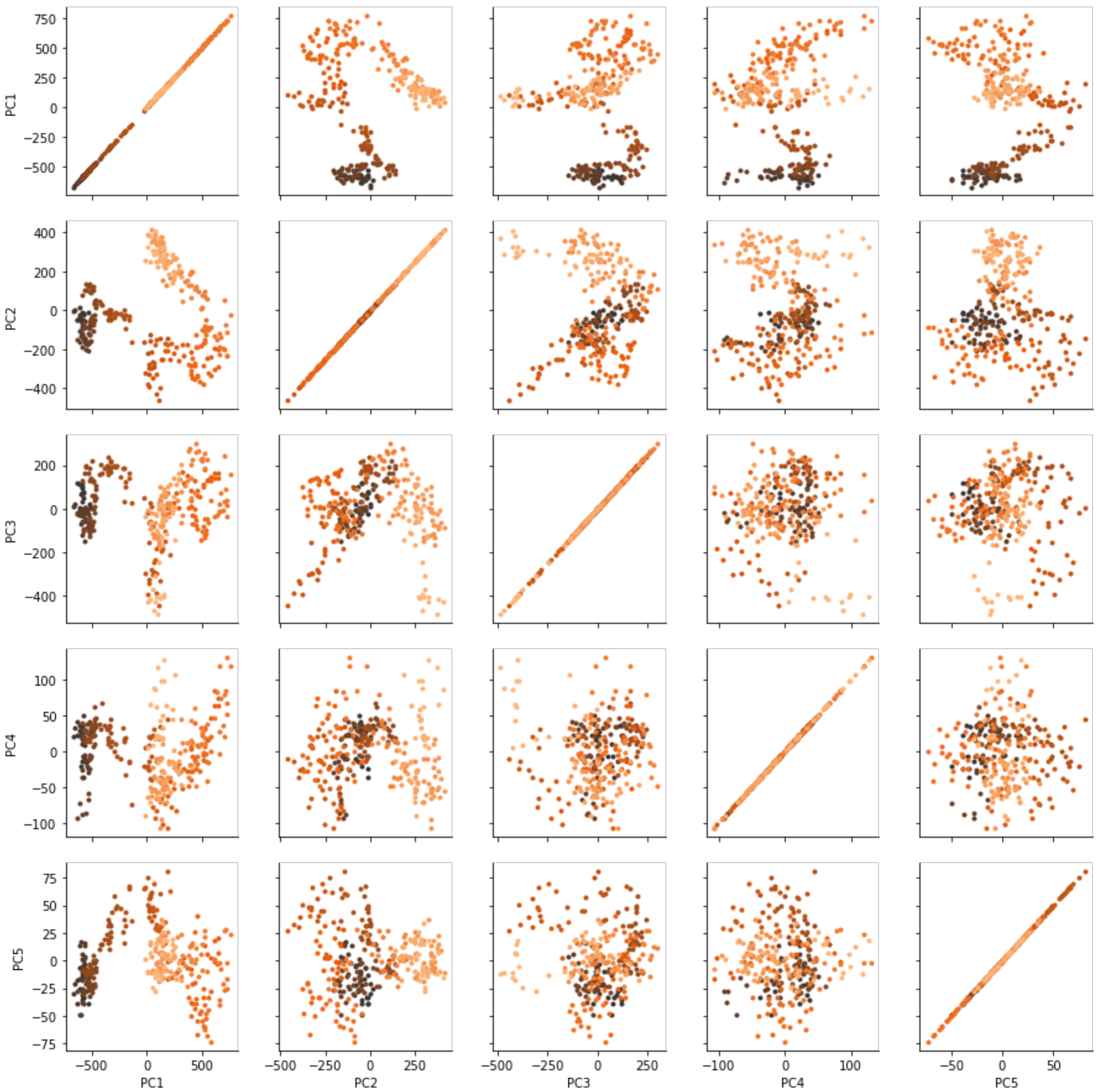

**Figure S12.** Principal component pair-wise interaction map for A1 variant system. First 5 principal components are shown. Color corresponds to time and goes black to dark orange.

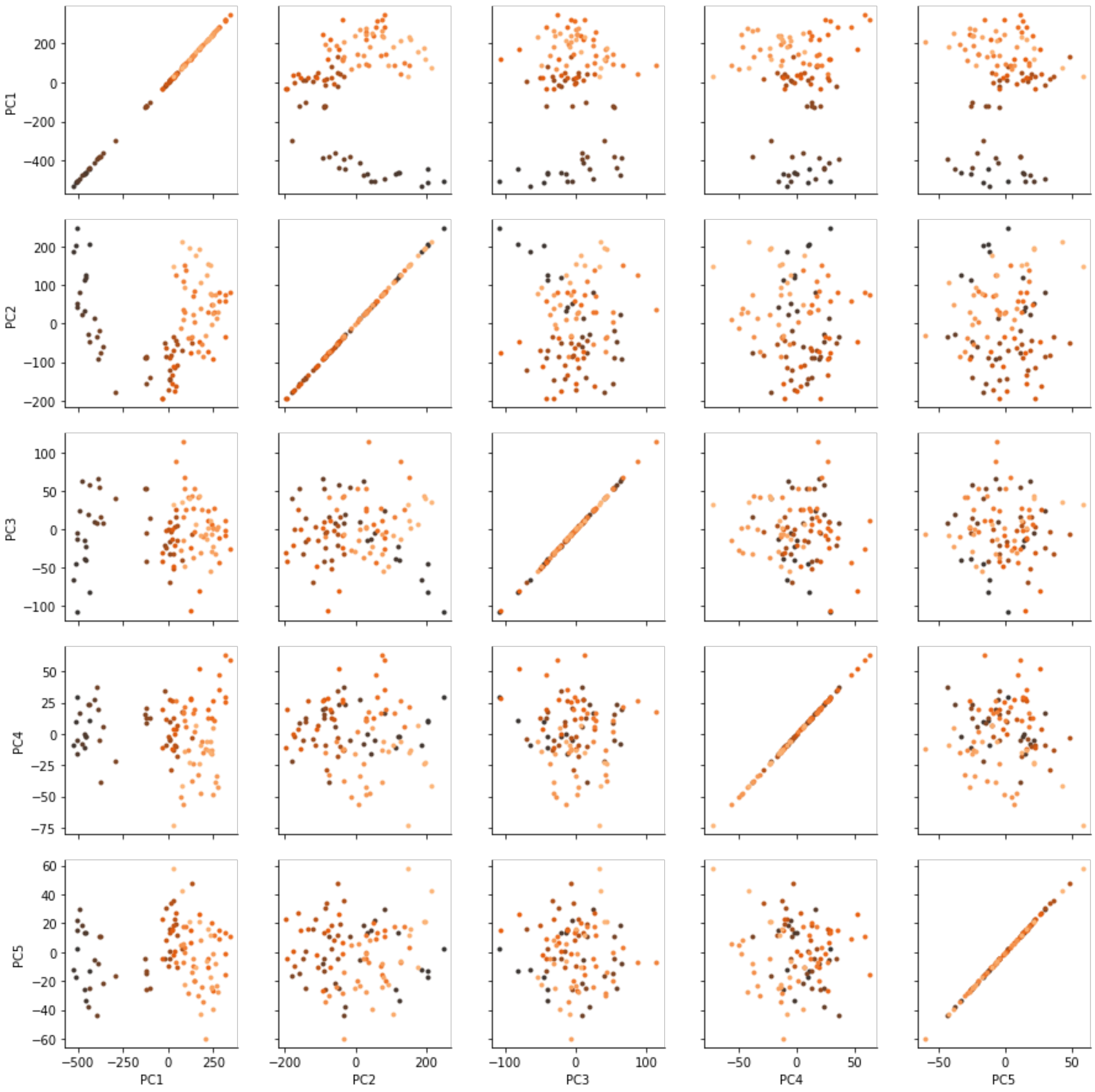

**Figure S13.** Principal component pair-wise interaction map for A2 variant system. First 5 principal components are shown. Color corresponds to time and goes black to dark orange.

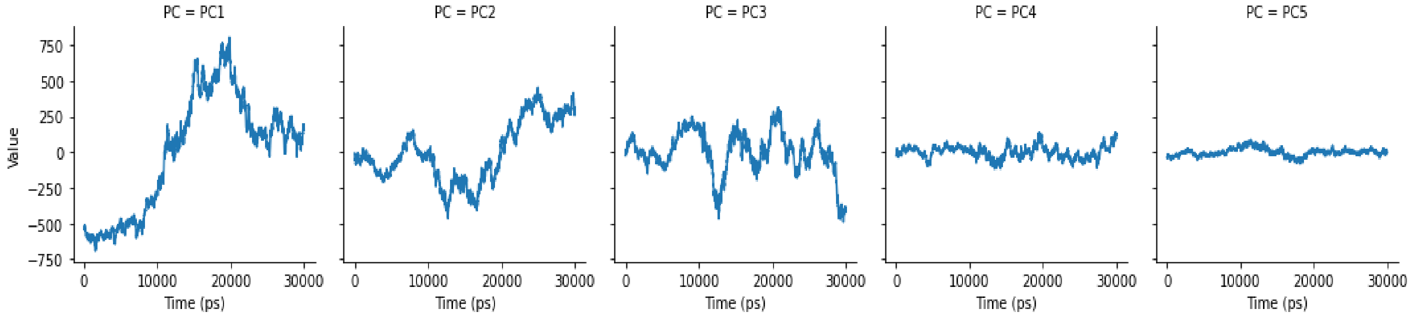

**Figure S14.** Principal component vs time for A1 variant system.

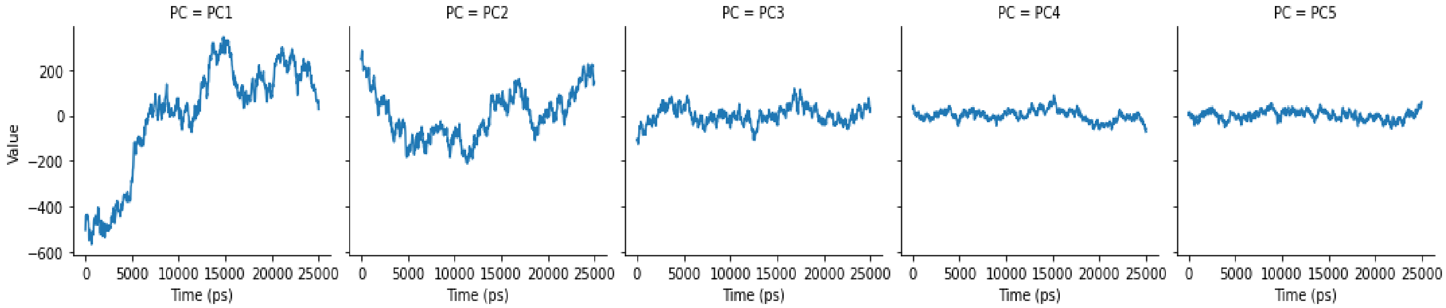

**Figure S15.** Principal component vs time for A2 variant system.
